# From cells to tissue: How cell scale heterogeneity impacts glioblastoma growth and treatment response

**DOI:** 10.1101/650150

**Authors:** Jill A. Gallaher, Susan C. Massey, Andrea Hawkins-Daarud, Sonal S. Noticewala, Russell C. Rockne, Sandra K. Johnston, Luis Gonzalez-Cuyar, Joseph Juliano, Orlando Gil, Kristin R. Swanson, Peter Canoll, Alexander R. A. Anderson

## Abstract

Glioblastomas are aggressive primary brain tumors known for their inter- and intratumor heterogeneity. This disease is uniformly fatal, with intratumor heterogeneity the major reason for treatment failure and recurrence. Just like the nature vs nurture debate, heterogeneity can arise from heritable or environmental influences. Whilst it is impossible to clinically separate observed behavior of cells from their environmental context, using a mathematical framework combined with multiscale data gives us insight into the relative roles of variation from inherited and environmental sources.

To better understand the implications of intratumor heterogeneity on therapeutic outcomes, we created a hybrid agent-based mathematical model that captures both the overall tumor kinetics and the individual cellular behavior. We track single cells as agents, cell density on a coarser scale, and growth factor diffusion and dynamics on a finer scale over time and space. Our model parameters were fit utilizing serial MRI imaging and cell tracking data from ex vivo tissue slices acquired from a growth-factor driven glioblastoma murine model.

When fitting our model to serial imaging only, there was a spectrum of equally-good parameter fits corresponding to a wide range of phenotypic behaviors. This wide spectrum of *in silico* tumors also had a wide variety of responses to an application of an antiproliferative treatment. Recurrent tumors were generally less proliferative than pre-treatment tumors as measured via the model simulations and validated from human GBM patient histology. When fitting our model using imaging and cell scale data, we determined that heritable heterogeneity is required to capture the observed migration behavior. Further, we found that all tumors increased in size after an anti-migratory treatment, and some tumors were larger after a combination treatment than with an anti-proliferative treatment alone. Together our results emphasize the need to understand the underlying phenotypes and tumor heterogeneity in designing therapeutic regimens.

## I. Introduction

Glioblastoma (GBM) is the most common and deadly form of brain cancer with a median survival rate of 12-15 months (1,2). The extensive infiltration of single cells in and around important anatomical structures makes curative surgical resection practically impossible, and resistance to radiation and chemotherapeutic strategies often causes recurrence following an initial response. Magnetic resonance imaging (MRI) serves as the primary diagnostic viewpoint into the disease state and guides the subsequent treatment strategies that follow. However, it is often the case that patients with similar growth patterns determined with MRI will have different post-treatment kinetics. While patient data at smaller scales, such as histological and genetic profiling, is known to be generally prognostic, its connection to optimal therapeutics and clinical imaging remains an active area of research (3–8). In this work, we investigate how phenotypic heterogeneity at the cell scale effects tumor growth and treatment response at the imaging scale by quantitatively matching multiscale data from an experimental rat model of GBM to a mechanistic computational model.

It is broadly acknowledged that GBMs exhibit genetic and phenotypic heterogeneity both spatially and temporally (9–13). However, GBM progression is not just driven by cell autonomous genetic and epigenetic alterations but also from larger scale non cell autonomous interactions between cells and their environment (14–16). Data is routinely collected in the clinic, but different scales are generally separated. Imaging gives us larger tissue scale information like size to quantify burden or density variations that can be used to define different environmental habitats (17–19). Histology, single cell data, and genetic profiling can be used to view heterogeneity at the tissue and individual cell level, however, the measured heterogeneity at the cell scale does not directly lead to predictions in tumor growth and treatment response.

Here we examine feedback between tumor and microenvironmental heterogeneity using a model that considers amplification of platelet-derived growth factor (PDGF). PDGF is a potent mitogen that appears to be important for invasion and expansion of proneural GBM (14,20–27). PDGF can stimulate proliferation, migration, and differentiation of normal progenitor cells (28,29) and tumor cells (30). Cells may encounter different local external PDGF signals and also have a variable response to PDGF. While a transient PDGF signal is part of a normal injury repair response mechanism (28,29), glioblastoma tumor cells can also overexpress PDGF to drive tumor growth. Whilst it is impossible to separate observed cell phenotypes from their environmental context *in vivo*, we can investigate this complex system using a mathematical framework coupled to multiscale data to get a more complete picture of the disease (Fig. 1). In this work, we use MRI imaging data and *ex vivo* time lapse imaging of fluorescently tagged cells in tissue slices (Fig. 1 upper) to parameterize a mechanistic hybrid agent-based model (Fig. 1 lower).

**Figure 1.**
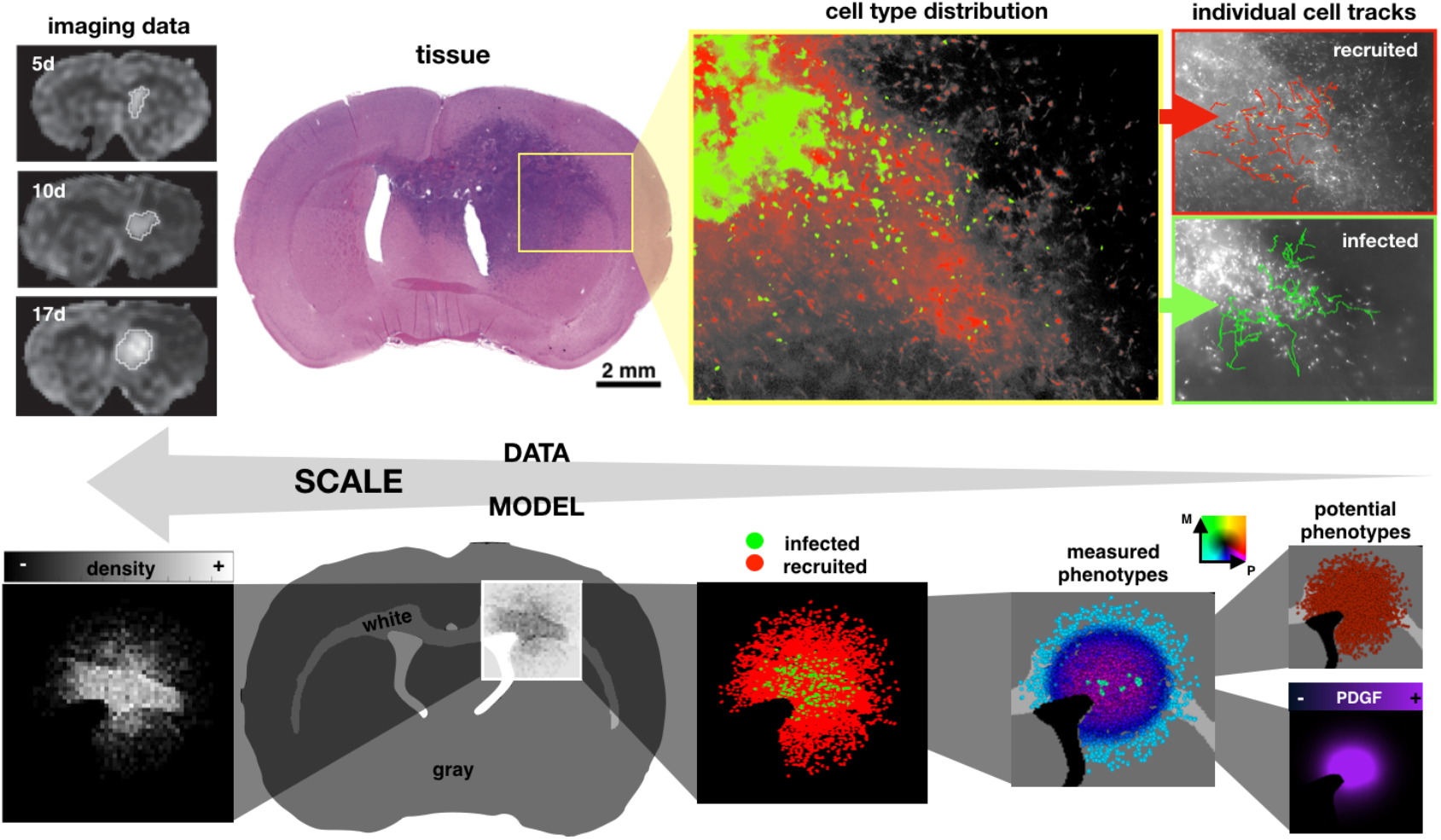
Coupling multiscale data to a multiscale mathematical model. Upper: data from rat experiments including imaging at 5, 10, and 17 days post injection, circumscribed and quantified from serial MRI images, tissue slice image, spatial distribution of infected (green) and recruited (red) cells, and individual cell tracks. Lower: the multiscale model represents the imaging as a spatial density map, considers the gray and white matter distribution in the rat brain tissue, and tracks cell types (infected and recruited), measured cell phenotypes (actual proliferation and migration), potential cell phenotypes (maximal proliferation and migration), and the PDGF concentration field.

Mathematical models have been developed to study many facets of GBM growth and response to treatment (5,30–42). There have been numerous papers published by Swanson *et al* demonstrating the clinical use of a relatively simple partial differential equation model based on net rates of proliferation and invasion. To date they have used their models to predict therapeutic benefits from surgery and radiation (43–46), IDH1 mutation status (47), and implications of growth kinetics during PDGF-driven tumor progression (33,34). However, the continuum nature of this model means it cannot capture intercellular heterogeneity which may impact long-term post treatment behavior. Here, we consider intratumor heterogeneity in proliferation and migration rates from inheritable phenotypes at the cell scale and from the microenvironment. The multiscale nature of our hybrid model enables us to tune our parameters with both imaging and cell-tracking data, thus allowing us to predict a host of tumor behaviors from size to composition to individual cell responses to therapy. This could be key to understanding treatment response as single cells can cause relapse or treatment failure.

In the following sections, we introduce the experimental model by Assanah *et al* of PDGF-driven GBM in which single cells were tracked. We then present a hybrid agent-based mathematical model which is able to capture the spatial and temporal heterogeneity of single cells. Using this model, we first identify the sets of parameters with which our model is able to recapitulate the observed tumor size dynamics from the data. We then identify the sets of parameters that fit smaller scale metrics from the data, such as the observed distribution of individual cell velocities. We investigate how the fully parametrized model with both environmental and heritable heterogeneity compares to a case with only environmental heterogeneity, and finally, we show how anti-proliferative and anti-migratory drugs affect outcomes and modulate heterogeneity within the tumor cell population.

## II. Methods

### Rat model and *ex vivo* multiscale data analysis

The experimental rat model enabled the tracking of both cells that were infected with the PDGF-over-expressing retrovirus, tagged with green fluorescence protein (GFP), and normal progenitor cells, tagged with dsRed. At 2 and 10 days post infection, brains were excised and cut into 300μm thick slices, and positions of labeled cells and their progeny were tracked by hand every 3 minutes from time-lapse tracking. For more details on the experimental model, see (14). A total of 611 cells were tracked (134 infected and 137 recruited at 2d and 137 infected and 203 recruited at 10d) in the tissue slices (2 slices at 2d and 4 at 10d) over time. Proliferation rate was calculated by dividing the number of proliferation events over the time period by the total number of cells at the beginning of the observation period and the total observation time in hours. For each cell we calculated a cell speed by the total distance traveled over the total time spent moving. The persistence times for moving and stopping, and the turning angles were also calculated (see Section S1).

### Hybrid off-lattice agent-based mathematical model

Our hybrid model consists of tumor cells, represented as off-lattice agents, and a PDGF distribution, represented as a continuous field. We used off-lattice agents to allow single cells to migrate without the confines of a grid structure, but used a larger scale square lattice to track the cell density matrix, which we used to check if the local carrying capacity was reached. A smaller hexagonal lattice was used to track PDGF dynamics and define the brain tissue in terms of white and gray matter.

#### Model initialization and flow

We define the white and gray matter using a section from an 80 day old male Sprague Dawley rat (48–50) using the Scalable Brain Atlas (51). We selected a coronal slice near the bregma to get a representative 2D brain field involving the corpus callosum (Fig. 1 bottom). For simplicity, any anatomical tissue feature that was not white matter was rendered as gray matter. The final array defines an 833×573 pixel domain corresponding to a scaled brain size of roughly 14.5×10.0 mm. There is an initial injection of 100 infected cells, which are labeled green and produce PDGF, and 100 progenitor cells, which are labeled red and do not produce PDGF. In addition, glial progenitors are randomly initialized throughout the brain matter at variable density around 2% (52,53), and there is an initial bolus of PDGF, representing an injury response caused by the injection (14). The flowchart in Fig. 2A details the major decisions at each time point about division (orange), migration (teal), and PDGF (purple). All cells are assumed to be 25 μm in diameter.

**Figure 2.**
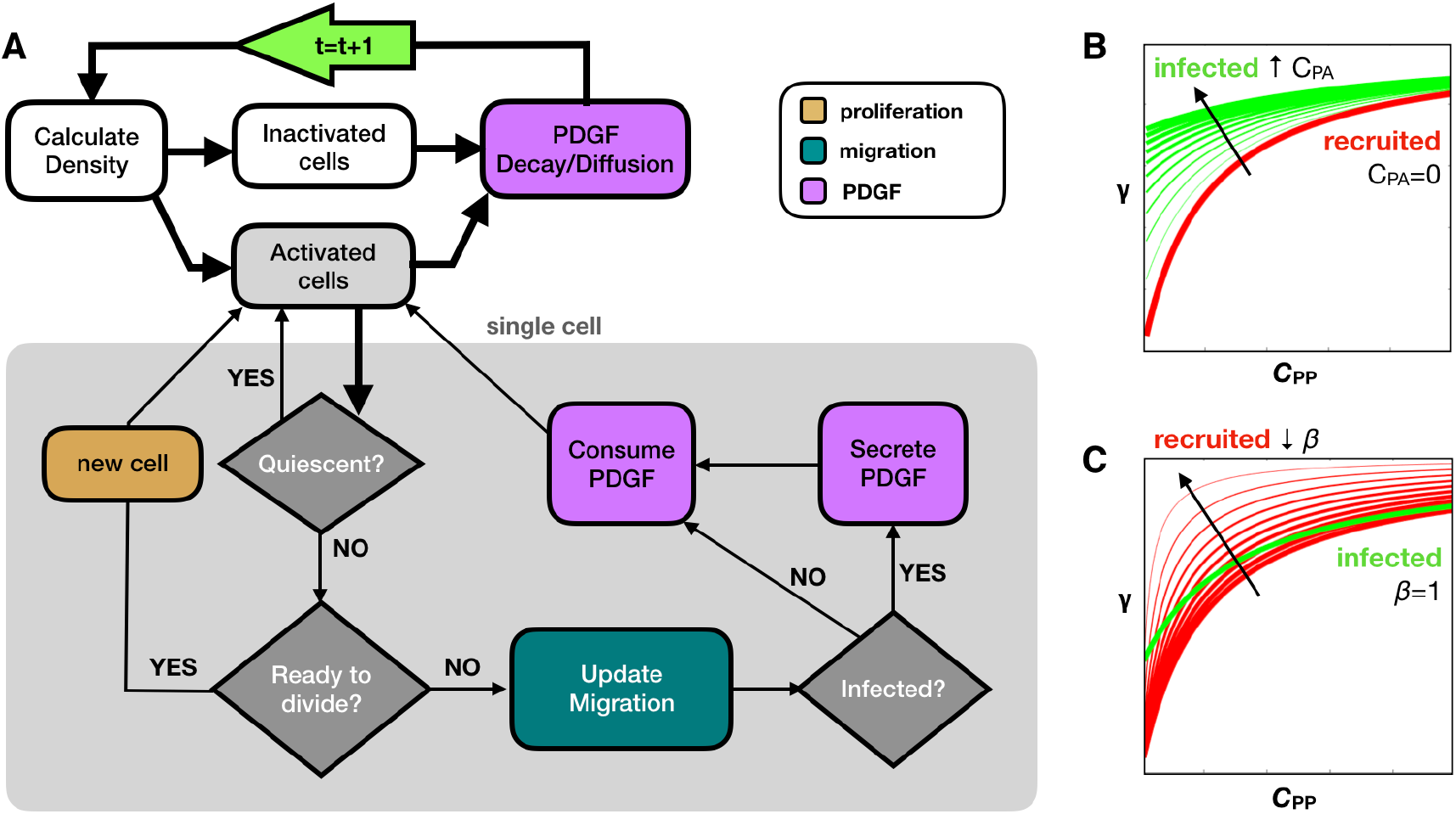
Model overview. A) Flow chart shows key decision points in the model. Tissue processes are connected with thick black lines, while the cell loop for single cell processes are contained within the gray box and connected with thin black lines. At the start of each time step (green arrow), we calculate the density and find the activated and inactivated subsets of cells. All activated cells are checked for quiescence, division, migration, and PDGF interactions as shown. Then PDGF decay and diffusion occurs before moving onto the next time step. The infected and recruited cells respond differently to PDGF due to B) an autocrine stimulation for infected cells (*C*_PA_ in Eq. 2) and C) a decreased activation barrier for recruited cells (*β* in Eq. 2). Increasing *C*_PA_ shifts the response upward at low *C*_PP_. Decreasing *β* increases the slope to achieve high response at lower *C*_PP_, while still inactive at *C*_PP_=0.

#### Calculate cell density matrix

We define a coarse square mesh (100μm × 100μm) to check the local cell density. Each cell is assigned a closest neighborhood, which has a carrying capacity of κ in gray matter and 2κ/3 in white matter. We also check progenitor cell activation at this step, as only activated cells go through the cell loop. The field of progenitor cells remain inactive unless the local PDGF is greater than 5×10^−4^ng/mL.

#### Cell loop

##### A. Proliferation and quiescence

A cell’s intermitotic time acts as a timer for division, counting down at each time step until the end of the cycle. At that point, a new cell is created at a random angle one radius away from the parent cell. However, if the number of cells in the neighborhood mesh point exceed the carrying capacity, then it is deemed quiescent, and it does not move forward in its cell cycle and does not divide. If subsequently there is enough room to divide, the cell reenters the cell cycle where it left off. The newly divided cell inherits the same proliferation rate and migration speed as its parental cell.

##### B. Migration

Glioma cells migrate in a stop and go fashion (54). We randomly choose a migration status (stop or go), and sample from the distribution of persistence times. For a given persistence τ, a stopped cell will remain stopped and a moving cell will continue to move at the current angle and velocity. After τ time, the cell resets its migration status (stop or move), resamples τ from the data, and finds a new moving angle. In gray matter, cells do a random walk for τ sampling from a uniform distribution of turning angles, and in white matter, cells do a persistent random walk for 1.5τ sampling from a normal distribution centered around 0 with a standard deviation θ. A cell is not allowed to move into empty space, such as past the edges of the brain or within the ventricles. If a cell lands in this space, it has 10 attempts to find a suitable spot at other random angles. If unsuccessful, the distance moved is increased by a cell diameter, and the angle search is repeated for distances of up to 3 diameters away from the original location. If an empty space is not found, the cell remains in the original location (however, in our testing, a new location was always found before this constraint was satisfied). If the cell is set to move into a space that is already at carrying capacity, then it can move there only if it is less dense than the original space. Otherwise, it remains in place. This allows the density of cells to slightly surpass the carrying capacity but prevents much movement when above or near the carrying capacity.

##### C. Response to PDGF

PDGF can stimulate glial cells to proliferate and migrate by autocrine and paracrine signals (28,55). Since we are interested in phenotypic heterogeneity in the tumor with regards to proliferation and migration, we need to separate the influence of the environmental PDGF, which can change depending on location, from the potential phenotype, which is inherited. To achieve this, we model the cells such that their observed phenotype for proliferation rate *p* and migration speed *m* is a product of the response to PDGF in the environment and some internal, inheritable upper limit:

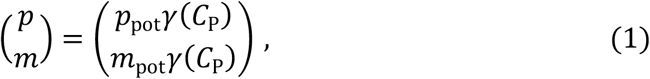

where *p*_*pot*_ is the maximal potential proliferation rate, and *m*_*pot*_ is the maximal potential migration rate. The function *γ*(*C*_P_) represents how the concentration of PDGF *C*_P_ modulates the proliferation and migration, which ultimately takes a value from 0-1, so that *as C*_P_ becomes saturated proliferation and migration reach their maximum potential values (i.e. *γ(C*_P_)→1, so that *p*→*p*_*pot*_ and *m*→*m*_*pot*_). The exact functional relationship of *C*_P_ on *p* and *m* is not well established, but a Hill function response in compatible with the data (30,56):

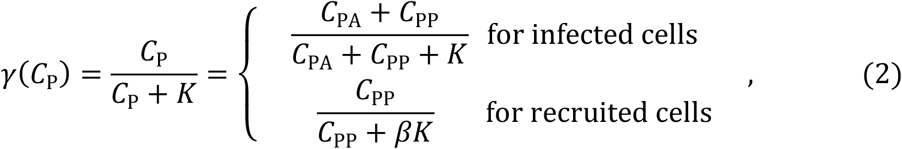

where *C*_PA_ is the PDGF contributing to the autocrine stimulation, *C*_PP_ is the PDGF contributing to the paracrine stimulation, K is the concentration at which the response is half maximum, and *β* modifies the activation barrier of recruited cells to PDGF stimulation. While all cells can respond to PDGF produced by the infected cells that diffuses throughout the surrounding environment *C*_PP_, only the infected cells have an autocrine effect, due to a portion of the PDGF *C*_PA_ that stays within and stimulates the infected cells. The recruited cells are also assumed to have a lowered activation barrier to *C*_PP_. We incorporate this into the equation by lowering the concentration at which the response is half maximum (by modifying *K* by *β*∊(0.1,1)), which causes recruited cells to gain a larger response from *C*_PP_ than infected cells while still being inactive when *C*_PP_=0. The effects of changing these values are shown in Figs. 2C-D. Because there are a large number of inactive recruited cells in the environment, we cut off any activity from these cells in areas with *C*_PP_<5×10^−4^ng/mL, which corresponds to an upper bound of 0.1% for the response function (*γ*(*C*_P_)<=0.001) with the given parameter ranges. This cutoff reduces the computational expense from behavior that is essentially negligible.

##### D. PDGF secretion and consumption

Only infected cells secrete PDGF and all cells consume PDGF into or from the nearest hexagonal grid point. If there is less local PDGF than the amount to be consumed for a cell during the time step, all PDGF in the grid point will be consumed.

#### PDGF dynamics

A fine hexagonal mesh with the same radius of a cell (12.5 μm) is utilized for the PDGF dynamics. Following the cell loop, the whole PDGF field is subject to decay and then diffusion (further details in Section S2).

## III. Results

### Cell behavior in *ex-vivo* assay is influenced by multifaceted factors

In a series of experiments by Assanah *et al*, it was shown that infecting resident glial progenitor cells with a retrovirus engineered to overexpress PDGF in the rat brain can induce a massive overgrowth of cells with histologic features similar to GBM (14,22). The tumors grow rapidly and are composed of a mixture of retrovirus infected and uninfected/recruited progenitor cells (14). Specifically, the tumor diameters at 5, 10, and 17 days post infection were 1.7, 2.4, and 3.2 mm, respectively, which were determined previously from MRI images in Massey *et al* (33). At 17d, progenitors made up 80% of all labeled cells in the tissue section (14). Single cell trajectories from the infected (green) and recruited (red) cells at 2d were tracked and are displayed in the spatial plot of Fig. 3 along with births, stops, and speeds along the tracks. Cells were mainly measured near the edge of the tumor where the density was lower, so they could be distinguished from their neighbors. We found that there was a high degree of phenotypic heterogeneity amongst cells, some of which may be due to environmental influences. This is outlined below.

**Figure 3.**
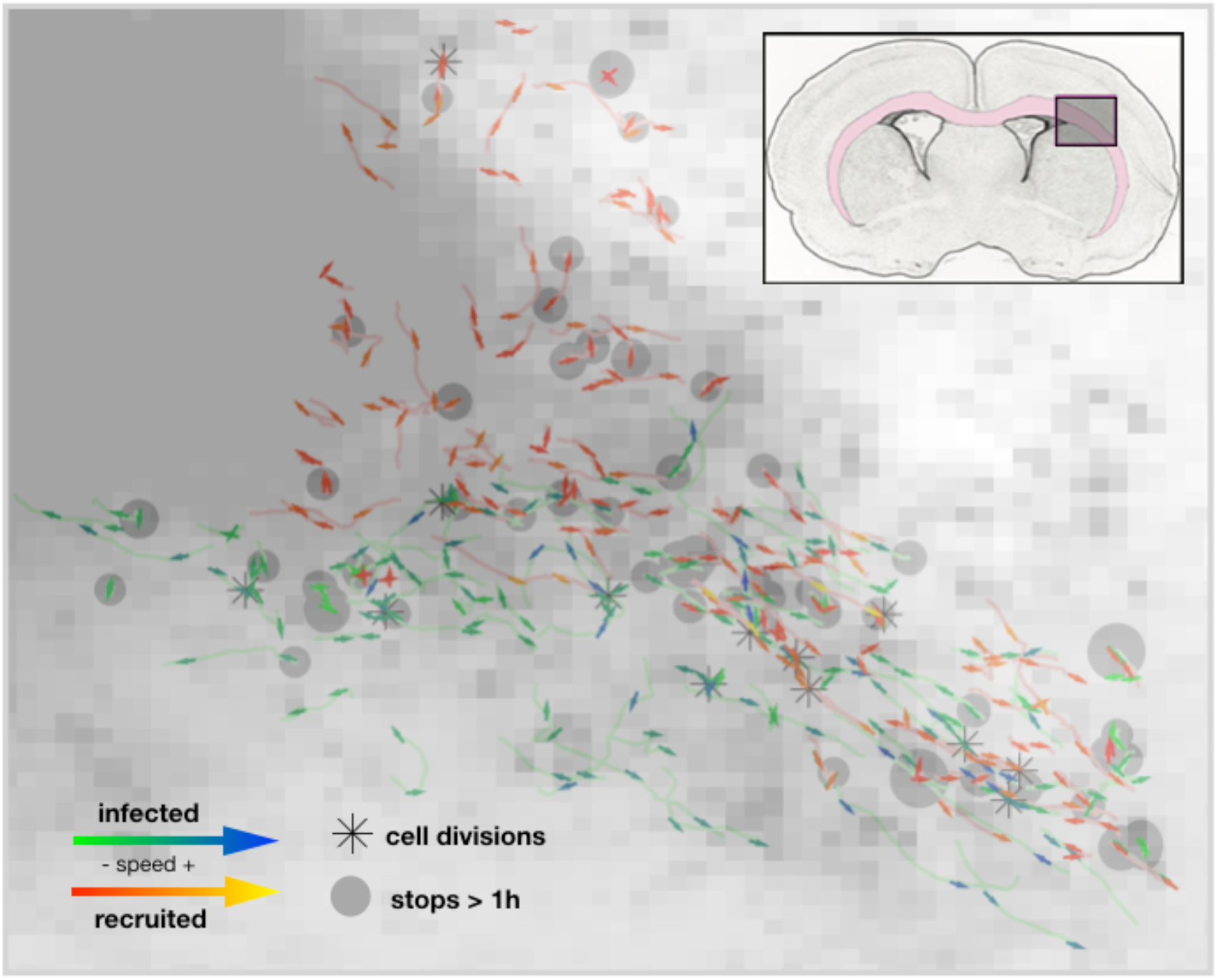
Single cell trajectories from the rat experiment at 2 days post infection overlaid on the cell density map. The insert shows the region of interest within the rat brain where the pink highlights the white matter. An asterisk marks where a cell division occurred. Each track contains an arrow for the first and last half of the track showing the average direction and speed over that time period. The arrows for the infected cells are green for lower speeds and blue for higher speeds. The arrows for recruited cells are red for lower speeds and yellow for higher speeds. Gray dots mark where a cell has stopped longer than 1 hour with the size proportional to the stop time.

#### Phenotypic heterogeneity

From these tracks, we were able to observe where cells moved, divided, turned, and stopped for long periods of time. They generally moved in the same direction, but occasionally made large turns and took long stops. There was large variation in the speeds of the cells. The average speed was slightly higher for recruited cells, but didn’t differ much between the different time points. The long stops and the cell divisions were scattered throughout the tissue and didn’t significantly correlate to the local density or each other. About half of the cells divided over the 25h track recording at 10d, and no cell during this time period divided twice. Proliferation rate was quantified as the percentage of cells that divided over time, which increased from 2d to 10d and was slightly higher for recruited cells (in agreement with the analysis in (14)). Plots of the migration behavior and a table quantifying the migration and proliferation metrics for this data from 2d and 10d are shown in Section S3.

#### Possible environmental influences

Cells appeared to move generally along the diagonal of the top-left to the bottom-right of the region, which corresponds roughly to the white matter region highlighted in pink in the insert of Fig. 3. There is also faster and more directional movement along the white matter tract while the denser areas of the tumor core and the outer gray matter areas generally had shorter, less directional paths.

### *In silico* tumors with similar growth dynamics may have widely different compositions

Using the multiscale data from the experimental model: tumor size over time, a count of cell types, and proliferation events and migration behavior tracked from single cells (Table S1), we calculate similar metrics in the *in silico* tumors (see Section S4A-B). We focused on a set of 16 uncertain parameter values with reasonably-defined search ranges (Table 1) and used a hybrid genetic algorithm-random sampling technique (57) to find parameter sets that fit the model to the time course of tumor sizes from the data at 5d, 10d, and 17d to within 10% error (Fig. S4).

**Table 1.** List of all variable trait ranges in the mathematical model. They are categorized into tissue-related, PDGF-related environmental effects, and cell specific values, such as response to PDGF or heterogeneity in proliferation and migration traits.

|  | PARAMETER | SYMBOL | RANGE (UNITS) | SOURCES |
| --- | --- | --- | --- | --- |
| tissue | Recruitable cell density | $\rho_R$ | 0.1-5 (%) | (52,53) |
| | directionality deviation in white matter | $\sigma_\theta$ | 0-45 (degrees) | - |
| PDGF | Initial PDGF | $p_0$ | 100-600 (ng/mL) | estimated |
| | Diffusion coefficient for PDGF | $D_p$ | 1-1000($\times 10^{-6}$ cm<sup>2</sup>/day) | estimated |
| | PDGF decay rate | $r_d$ | 0-0.500 (ng/mL.day) | estimated |
| | PDGF secretion rate | $r_s$ | 10-400 (ng/mL.cell.day) | (14) |
| | PDGF consumption rate | $r_c$ | (0-1)$r_s$ (ng/mL.cell.day) | - |
| PDGF response | Autocrine boost | $p_a$ | 0.1-50 (ng/mL) | (14) |
| | Half max proliferation response | $K_p$ | 5-300 (ng/mL) | (14,20,22) |
| | Half max migration response | $K_m$ | 5-300 (ng/mL) | (14,20,22) |
| | Recruited proliferation sensitivity | $\beta_p$ | 0.1-1.0 | - |
| | Recruited migration sensitivity | $\beta_m$ | 0.1-1.0 | - |
| proliferation | Intermitotic time ($p_{pot}^{-1}$) | $\tau$ | 20-100 (h) | (14,20,22) |
| | Std dev intermitotic time | $\sigma_\tau$ | 0-100 (h) | variable |
| migration | Migration speed ($m_{pot}$) | $v$ | 0-100 (μm/h) | (14,22) |
| | Std dev migration speed | $\sigma_v$ | 0-100 (μm/h) | variable |

The resulting tumors that fit the size dynamics encompass a broad range of distributions, shapes, and compositions. The results are shown in Fig. 4, with plots for metrics going from size dynamics to more smaller scale individual cell metrics (Fig. 4A-D). The diversity of best fits to the growth dynamics is plotted along with 3 examples that represent tumor densities that are more nodular (high density with a very distinct, steep border), diffuse (the tumor core is dense but drops off slowly in density), and intermediate. Spatial distributions for these 3 examples are shown at 17d. The size dynamics in Fig. 4A demonstrate that the best fits all have similar trajectories with little overall variation. However, the sizes in the simulation are determined by the average maximum diameter exceeding 10% of the carrying capacity. The many ways that the cells can be distributed and still meet the intended size to match the data are shown below Fig. 4A. The nodular tumor is relatively dense with a sharp drop at the edge, whilst the diffuse and intermediate tumors have more fuzzy borders due to a larger portion of cells distributed sparsely throughout the brain. These density differences can be quantified by defining respective tumor core diameters (at least 50% cell density) and rim sizes (tumor edge with at least 2% cell density). On average, the core diameters were 2.2mm, 1.9mm, and 1.9mm for the nodular, intermediate, and diffuse tumors, and the rim sizes were 0.4mm, 0.9mm, and 1.5mm respectively (Fig. S5).

**Figure 4.**
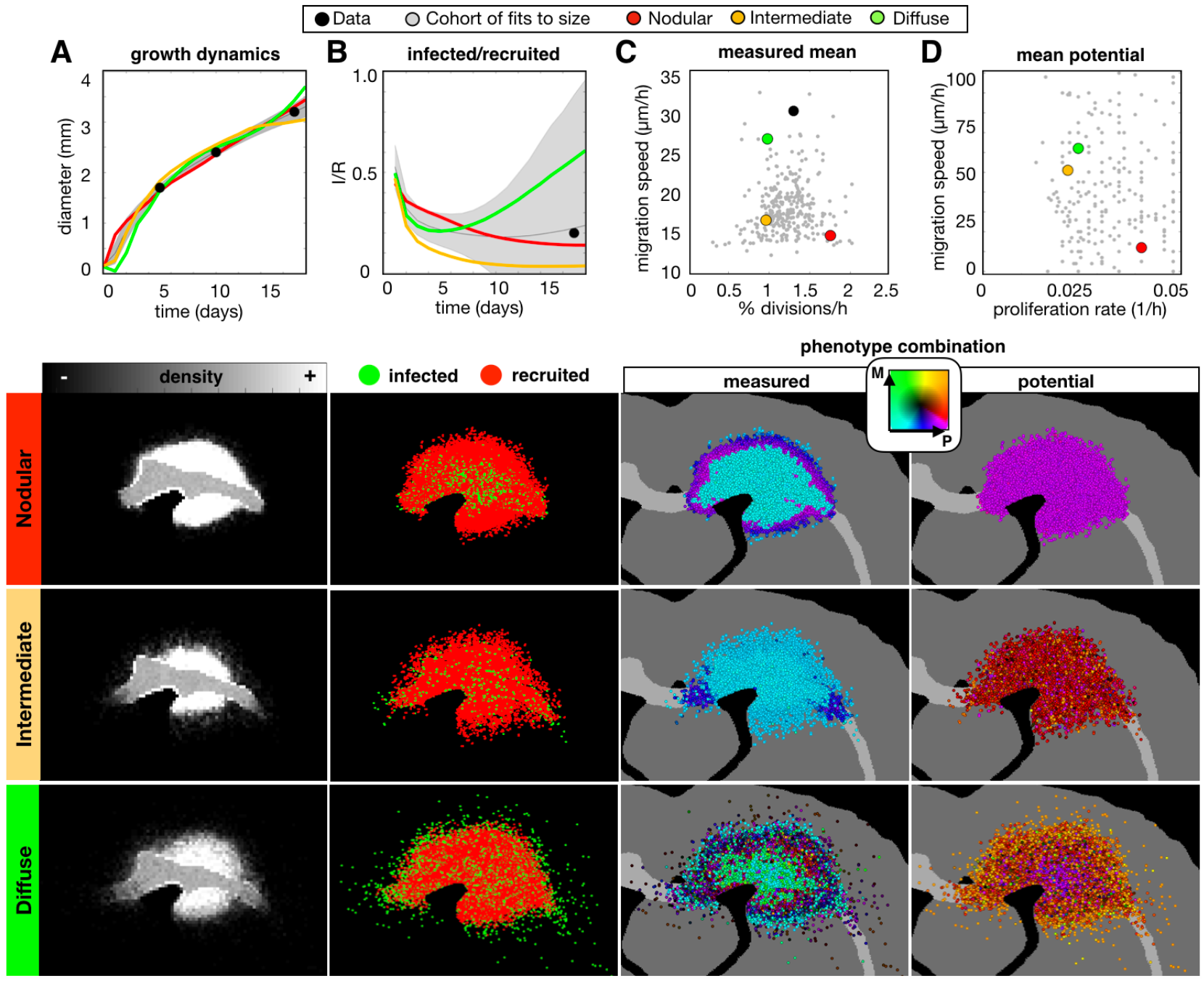
A wide range of *in-silico* tumors fit to the size dynamics from the experimental data. The top row shows the wider variation of the whole cohort of fits, while the spatial distributions below show representative nodular, diffuse, and intermediate density tumors at the 17d time point. The columns correspond to the (A) growth dynamics, (B) ratio of infected to recruited cells over time, (C) measured proliferation rate and migration speed averaged over all cells, and the (D) potential proliferation rate and migration speed (corresponds to the maximum values allowed given a saturated PDGF environment). For each metric, the data points are shown in black, the best fits to the size dynamics of the data are shown in gray (as a mean and standard deviation for dynamic values), and each example tumor is represented in the plots in color (as a mean over 10 runs). Parameter values for each tumor are given in Table S2.

While the size dynamics were similar amongst these tumors, smaller scale metrics differed substantially. Fig. 4B shows the variation in infected (I) and recruited (R) cell numbers. The nodular, intermediate, and diffuse tumors end up with I/R values of 0.17, 0.04, and 0.55, respectively. While both the nodular and intermediate tumors had more recruited cells along the periphery, the intermediate tumor had infected cells that extended farther along the white matter tracts. For the diffuse tumor, infected cells had advanced deep into the brain tissue in all directions.

The combination of measured trait values covered a large range of values (Fig. 4C). On average, the nodular tumor was more proliferative and less migratory, the diffuse tumor was more migratory and less proliferative, and the intermediate tumor had low values for both proliferation and migration. However, this differs spatially and is quantified over 10 runs for each tumor in Fig. S6A-B. High cell density, usually in the tumor core, creates a quiescent phenotype (characterized by suspended proliferation), which also varies amongst the tumors.

The potential phenotypes cannot be measured from the data but are of interest as they highlight difference between the realized (measured) and the possible (potential). The potential phenotypes are inherited over generations for each individual cell and represent maximal possible trait values. The nodular tumor is highly proliferative and minimally migratory throughout spatially and temporally. In contrast, the intermediate and migratory tumors are both initialized with similar potential phenotypes on average, however, they present as noticeably distinct tumors due to differences in heterogeneity and other parameter values. The effects of selection can be seen in the diffuse tumor, as the highly migratory and proliferative cells are found at the edge of the tumor and the less migratory cells are found in the tumor core. These effects are quantified in Fig. S6C-D.

### Anti-proliferative treatment causes a range of responses *in silico* tumors

We examined the effect of applying an anti-proliferative drug treatment, which represents a cytotoxic chemotherapy assumed to kill fast proliferating cells. We used a threshold cutoff of 60 hours, and all cells that are not currently quiescent with shorter intermitotic times than the threshold are killed. The drug was applied instantaneously at day 14 and remained on continuously until the simulation was stopped 28 days later. Figure 5 shows the results.

**Figure 5.**
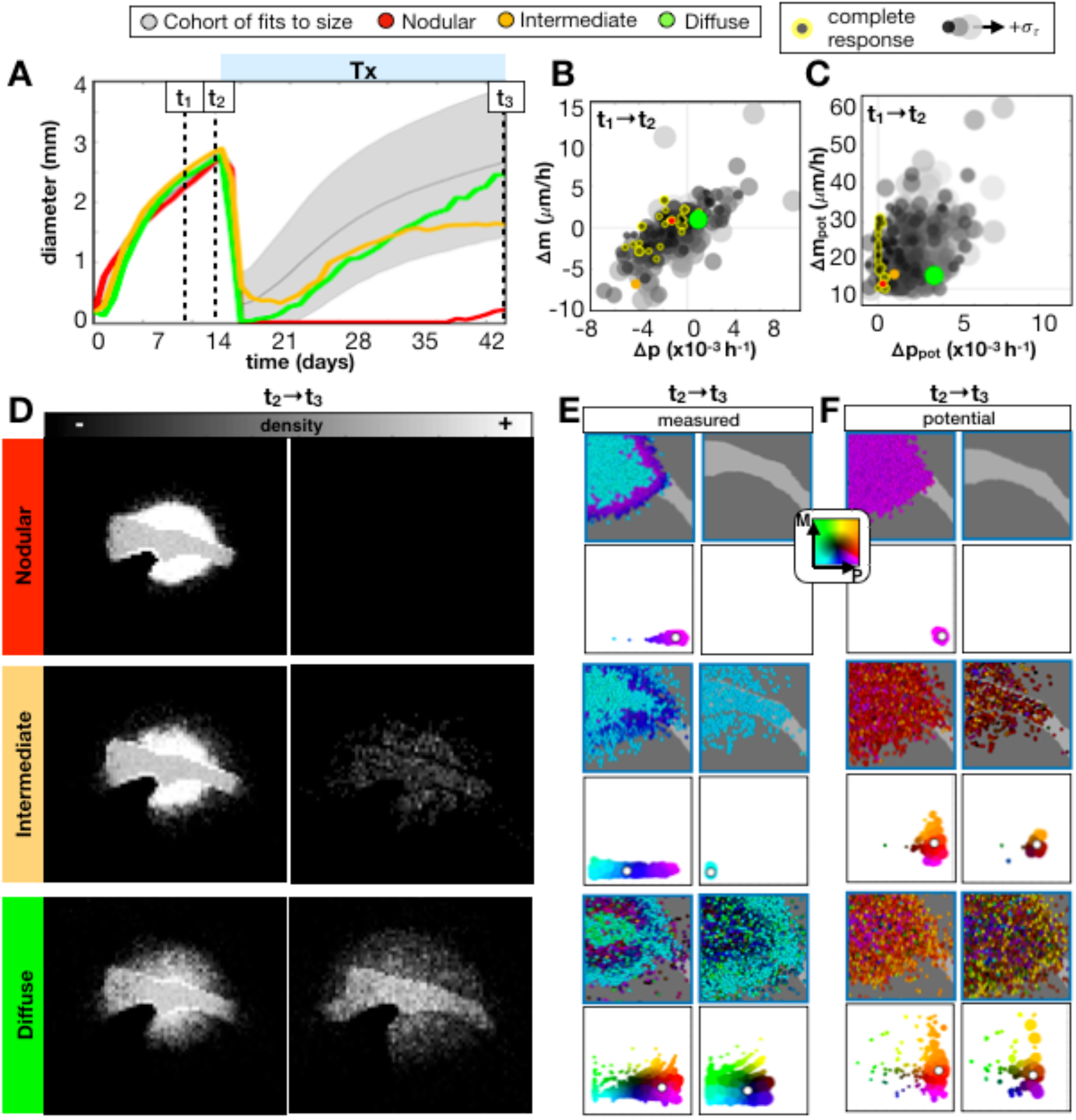
Range in long-term responses of *in-silico* tumors to an anti-proliferative drug. The drug was applied continuously at 14d until 42d. We compare the same top 300 fits and 3 tumors (averaged over 10 runs) shown in the previous figure. A) Growth dynamics before and during treatment. Change from t_1_=9d to t_2_=13d in B) measured and C) potential phenotypes. Larger, paler dots show tumors with more proliferative heterogeneity σ_τ_ and dots outlined in yellow represent complete responses to treatment. D) Density distributions of the nodular, intermediate, and diffuse tumors before (t_2_) and after (t_3_) treatment. The measured E) and potential F) phenotypes before (t_2_) and after (t_3_) treatment are shown spatially and as a scatter plot of phenotype combinations of only the non-quiescent cells). Both plots use the same color key, which depends on the proliferation rate and migration speed of the cells, but for the scatter plot the size of the circle is proportional to the number of cells with that phenotype combination, while a white dot marks the mean of the population.

Using the individual tumors examined in the previous section, we found that the nodular tumor most often showed a complete response to the anti-proliferative treatment, whilst the intermediate and diffuse tumors both recurred (Fig. 5A). Amongst all tumors in the cohort, there was a broad range of responses to the anti-proliferative treatment. In general, we found that the recurrent growth rate was the same or less than the pre-treatment growth rate.

We also examined whether any phenotype changes prior to treatment had predictive value. Within this cohort, a slowing proliferation rate was measured in tumors that had a complete response (Fig. 5B), but there was no significant trend in the measured migration rates. Generally, tumors had either decreasing activity in both proliferation and migration, no significant changes in either trait, or increased the activity of both proliferation and migration. The largest changes prior to treatment were observed from more heterogeneous tumors. Of the examples we examined in the previous section, the nodular and diffuse tumors did not change much over the observation period, but the intermediate tumor dramatically slowed in proliferation and migration, most likely due to recruitment of a large amount of progenitor cells.

While the measured proliferation rate was seen to slow prior to treatment for complete responses, we found that these tumors showed little to no change in the potential proliferation rate prior to treatment (Fig. 5C), but they were also rather homogeneous initially. Amongst the full cohort, there was a general trend toward both faster proliferation rate and migration speed that resulted in recurrence, and larger changes in the more initially heterogeneous tumors.

Changes to the representative nodular, intermediate and diffuse tumors are also noted post-treatment (Fig. 5D-F). The density distributions of recurrent tumors had similarly sized post-treatment images, but were more diffusely distributed than pre-treatment images (Figs. 5D and S7). The nodular tumor had very little variance in measured (Fig. 5E) and potential (Fig. 5F) phenotypes, and was rather homogeneously proliferative prior to treatment, while the two recurrent tumors had reduced mean proliferation rates upon recurrence. The intermediate tumor had less heterogeneity in both measured and potential traits upon recurrence compared to pre-treatment distributions, while the diffuse tumor remained heterogeneous.

### Cell autonomous heterogeneity causes little difference in tumor growth dynamics but can lead to big differences in response to treatment

To fit the model at the cell scale, we used the same parameter estimation method with all 16 measured observations from the experimental data. While the final best parameter set didn’t fit all metrics from the *in silico* model equally well to the data, the total error was within 15% for a cohort of parameter sets (Fig. S4C-D). Given the best fit parameter set from this group, we examined the effect of heterogeneity in the potential phenotype, such that eliminating heterogeneity would cause all observed heterogeneity to be environmentally driven, such as quiescence caused by high cell density and modulation of phenotype by local PDGF concentration. We compared the best fit parameter set (Heterogeneous) to one with the same mean potential values for proliferation and migration, but without heterogeneity in these rates, σ_*τ*_=0 and σ_*v*_=0, amongst the cells (Homogenous) along with the cohort of fits to all data within the 15% cutoff and the previous cohort of fits to the size dynamics alone (Fig. 6).

**Figure 6.**
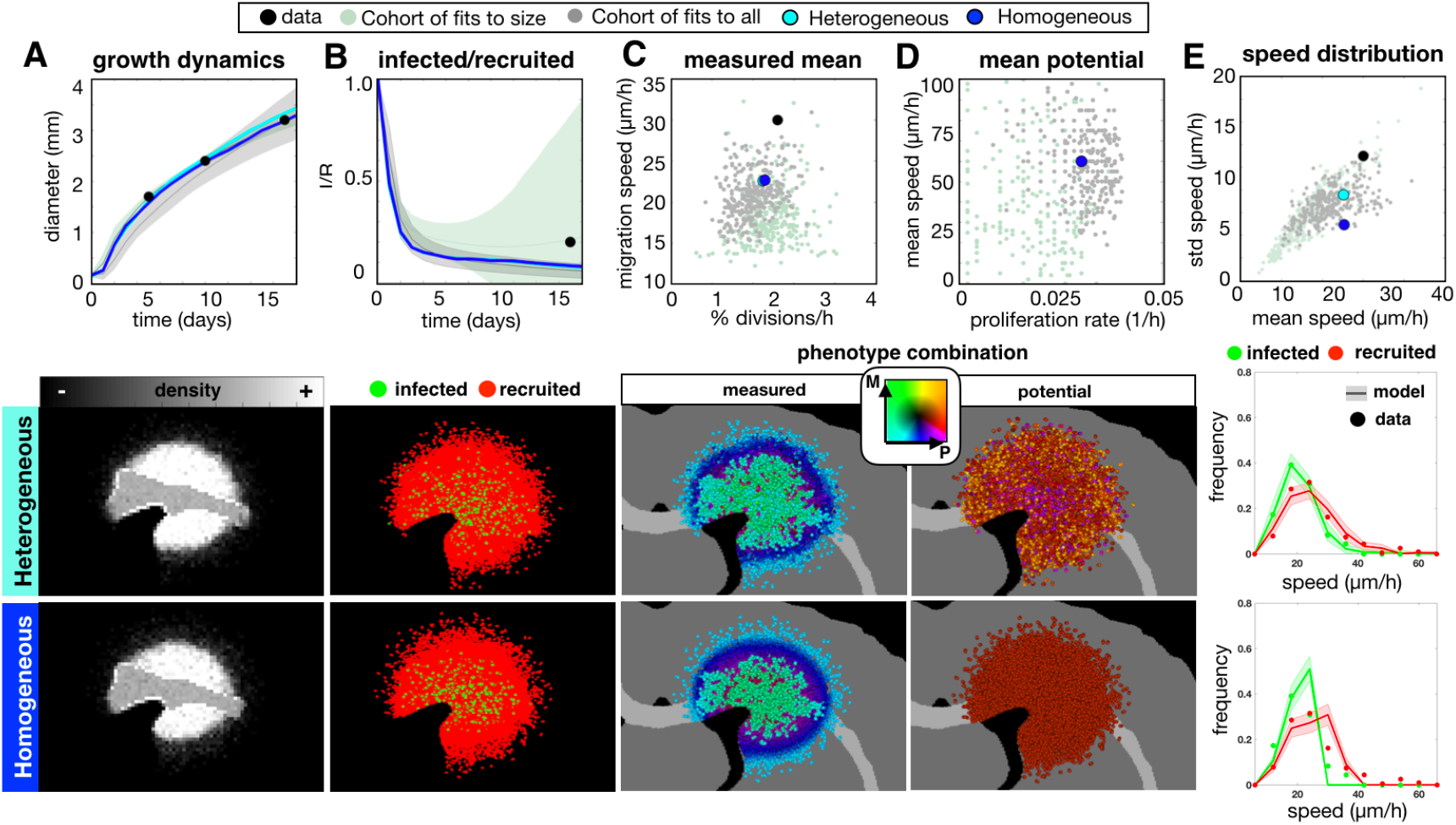
The top fit *in-silico* tumor to the multiscale experimental data using all 16 metrics. The top 300 fits to all data (gray) are compared to the top fits to just the size dynamics from Fig. 4 (green), the best Heterogeneous fit, and its Homogeneous counterpart (with no variation in potential phenotypes, i.e. *σ*_*τ*_=0, *σ*_*v*_=0). The data is in black. The For each metric, the corresponding spatial maps are shown below. Measured metrics include A) growth dynamics and B) infected/recruited cells over time, and at 10d the C) mean measured proliferation rate and migration speeds, the D) mean initial potential proliferation rate and migration speed, and the E) individual cell speed distributions in terms of mean and standard deviation. The final graphs in column E compare the 10d distributions of speeds of individual tracked cells to the data separated by cell type.

Fitting to all data, compared to just the size dynamics, proved to narrow the ranges to all metrics shown here with the exception of the size dynamics, which broadened slightly. Both the heterogeneous and homogeneous tumors reasonably fit the size dynamics (Fig. 6A) and had similar density distributions (Fig. S8). Both tumors and the larger cohort fit to all data underestimated the infected to recruited ratio (Fig. 6B). Both tumors had similar values for the measured proliferation and migration rates (Fig. 6C), showing that the observed heterogeneity is largely influenced by environmental drivers such as tumor density and PDGF concentration. Because the PDGF is highly concentrated at the tumor core and drops off at the tumor edge, the measured proliferation and migration rates reduce with the PDGF concentration (Fig. S9A-B). Both tumors were initialized with the same mean trait values (Fig. 6D), but the spatial distribution of potential trait values shows that heterogeneity in potential phenotypes can be present without manifesting any noticeable differences in larger scale metrics. For the heterogeneous tumor, effects of selection could be observed as the more migratory cells are found at the tumor periphery along with less proliferative cells (Fig. S9C-D). We also found differences in the distribution of individual cell speeds. The mean and standard deviation of speeds fit better when heterogeneity is present than when it is not (Fig. 6E), and comparing the distributions, which were averaged over 10 runs, further emphasizes this point (column 6E, lower). The *in silico* measurements for the heterogeneous tumor fit the data by not just matching to the peak, but also capturing the long tail of the distribution. The distribution for the homogeneous tumor drops off sharply at high cell speeds, which most likely occurs due to the maximum speed achieved at saturated PDGF levels. Only a small number of highly migratory cells like in the heterogeneous tumor is needed to create the long tail in this distribution.

When these tumors are treated with an anti-proliferative drug, there is enough heterogeneity in the heterogeneous tumor to cause recurrence and enough sensitivity in the homogeneous tumor for a complete response (Fig. 7A). The recurrent, heterogeneous tumor was slightly more diffuse on the edge (Fig. S10). There was a shift in both the observed (Fig. 7B) and potential phenotypes (Fig. 7C) prior to treatment to faster proliferation rates and faster migration speeds on average. The observed phenotypic activation increase is likely due to more sustained PDGF responses and heterogeneity necessary to fit the individual level metrics. Prior to treatment, the potential proliferation rates increased, while the potential migration speeds only slightly increased. On recurrence of the heterogeneous tumor, phenotypes with slower proliferation rates were, again, selected.

**Figure 7.**
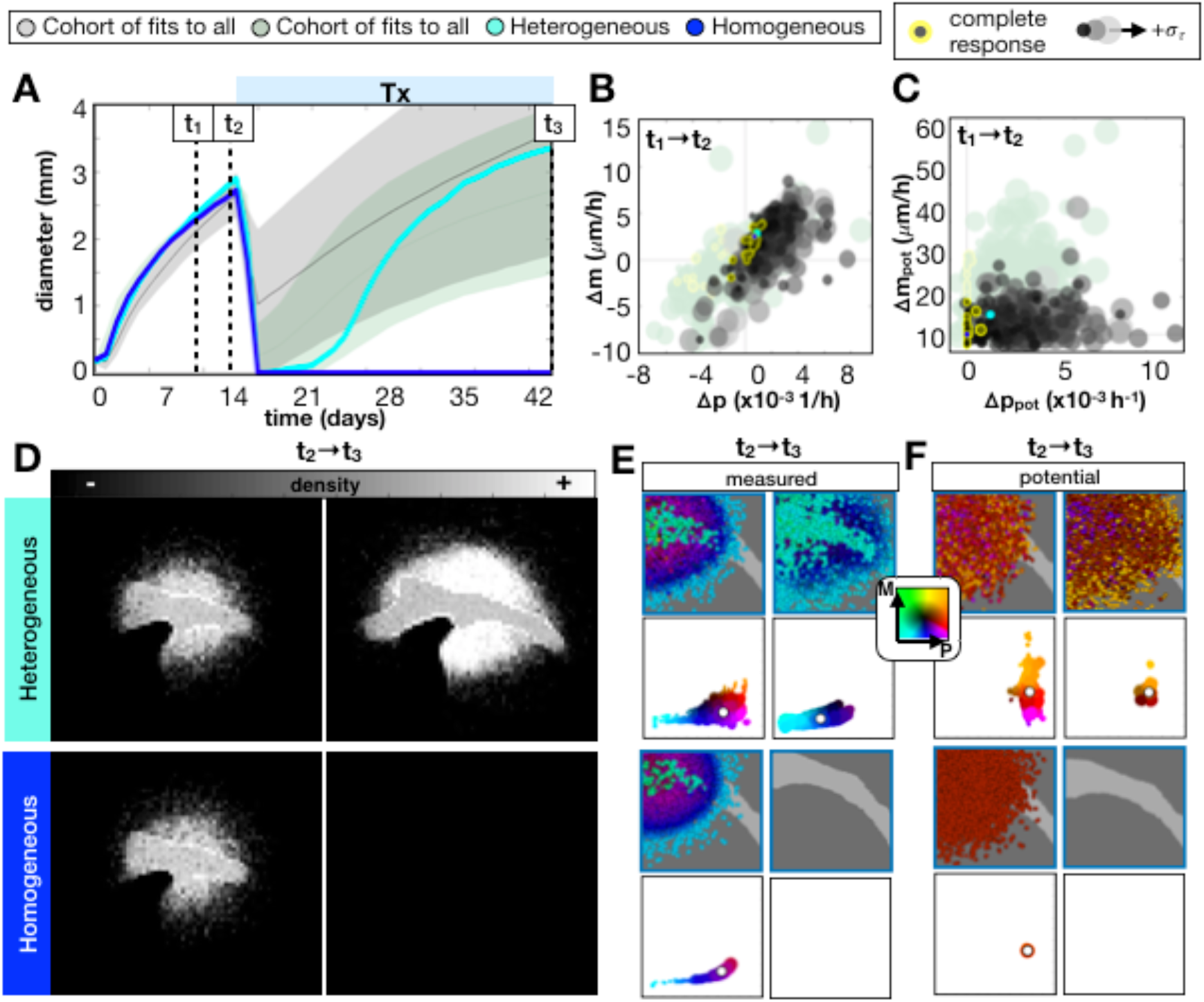
Range in long-term response of *in-silico* tumors to an anti-proliferative drug comparing fits to size vs. fits to all 16 metrics. The Heterogeneous best fit is compared to its Homogeneous counterpart (with no variation in potential phenotypes, i.e. σ_τ_=0, σ_v_=0). The drug is applied continuously at 14d until 42d. A) Growth dynamics before and during treatment are shown for the cohort of top 300 fits to all metrics, the previous cohort of 300 fits to sizes, and the heterogeneous and homogeneous tumors. Change from t_1_=9d to t_2_=13d in B) measured and C) potential proliferation rate and migration speed. The larger paler dots show tumors with less proliferative heterogeneity σ_τ_, and outlined in yellow are the tumors that showed a complete response to treatment. D) Density distribution of the heterogeneous and homogeneous tumors before (t_2_) and after (t_3_) treatment. The measured E) and potential F) phenotypes before (t_2_) and after (t_3_) treatment are shown spatially and as a scatter plot of phenotype combinations (of only the non-quiescent cells). The phenotype plots are represented as described in Fig. 5.

### Anti-proliferative treatment leads to a less proliferative tumor at recurrence in *in silico* and human tumors

Using the mathematical model, we found that antiproliferative drugs caused some degree of tumor recession over all cases tested, but the effect was often only temporary, and the recurring tumor had variable growth dynamics upon recurrence. Furthermore, there was some selection for slightly less proliferative cells, which give rise to recurrence. We also found similar results comparing the proliferating fraction of cells (Ki-67^+^) before and after chemoradiation for nine GBM patients (Fig. 8, upper). The proliferating fraction, measured through Ki67 staining, was seen to decrease upon tumor recurrence (p = 0.012, Wilcoxon matched-pairs signed rank test). In these cases, recurrence was defined as the first instance of measurable growth of the lesion on MRI with a clinical determination of disease progression resulting in a change of therapy, excluding pseudo-progression, in which the disease appears to progress and subsequently regress without change in treatment (59,60). Patients that demonstrated multifocal recurrence defined by multiple lesions not contiguous on MRI were excluded. Using a similar metric in the model to Ki67, we found similar results (Fig. 8, lower).

**Figure 8.**
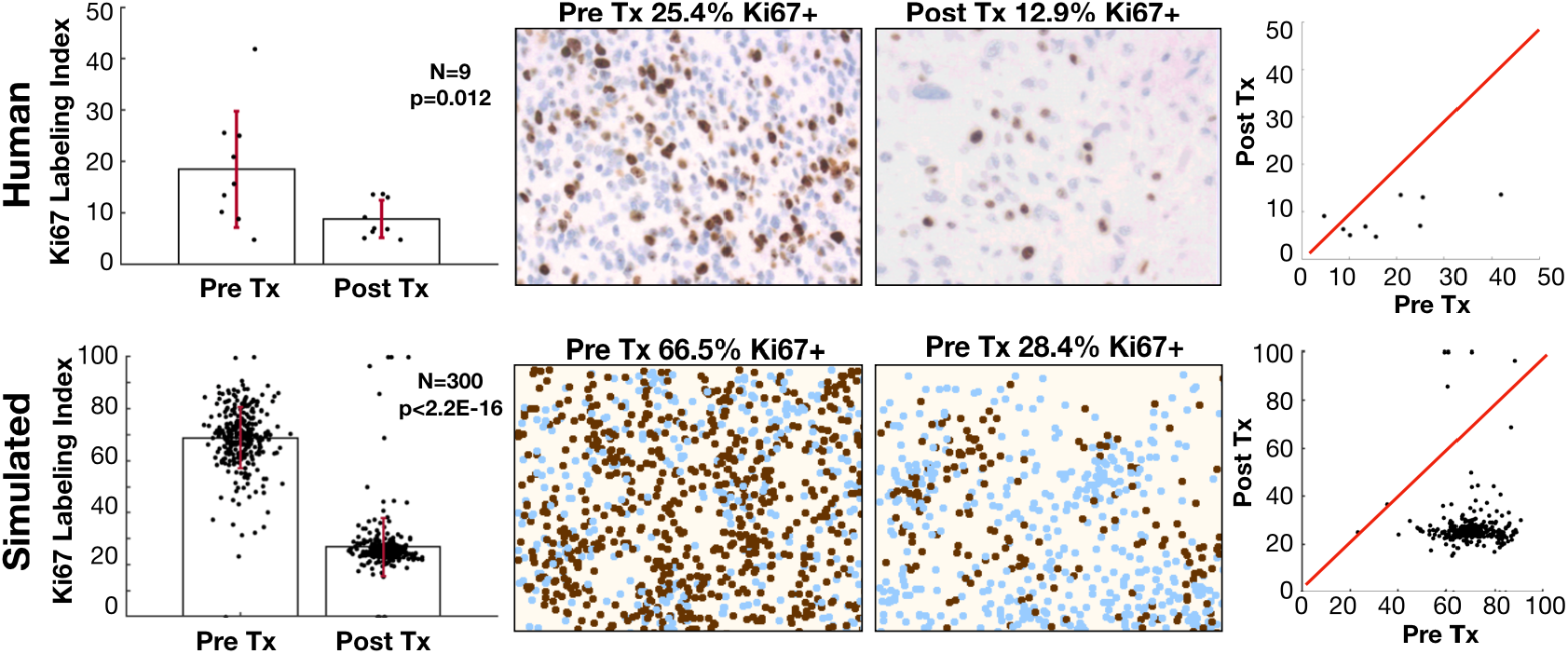
Proliferation is reduced in recurrent tumors. Upper: diagnosis and recurrent tumor specimens from 9 GBM patients stained with Ki-67 antibody indicating proliferating cells. Lower: pre-treatment (14d) and post-treatment (42d) proliferation index for the virtual cohort of fits to size dynamics. For the patient samples, the labeling index is defined as the % of DAB-stained area out of the total nuclear area for each patient in the region of highest staining density. For the model, we assume that Ki67 is positive only in the last 20 hours of the cell cycle, which is counted as a % in the area of highest activity. These are shown on the left with pre and post Tx variation and compared using a Wilcoxon matched-pairs signed rank test. The middle shows a representative pre and post Tx sample, and the right shows the correlation between pre and post Tx samples.

### Anti-migratory and anti-proliferative treatment combinations may improve outcomes in some *in silico* tumors

Anti-migratory drugs are an attractive option for very diffuse tumors to try to prevent further invasion into the brain tissue. We examined the effects of an anti-migratory treatment, represented as any agent that slows/stops the migration ability of cells (61,62). We simulated this treatment by slowing the migration speeds of all cells to 10% of their original speed. We compared an anti-proliferative treatment alone (AP), an anti-migratory treatment alone (AM), and an anti-proliferative and anti-migratory combination (AP+AM). We examined the effect of these treatments on the diffuse tumor from Fig. 4 as a prime example for an invasive tumor that could benefit from these treatments.

The *in silico* results show that the AM treatment alone is not successful in slowing the growth of most tumors, and the diffuse tumor grows especially fast under this treatment (Fig. 9A). Compared to the AP treatment, most *in silico* tumors do not do as well on AP+AM treatment at first, but appear to catch up over long applications of treatment.

**Figure 9.**
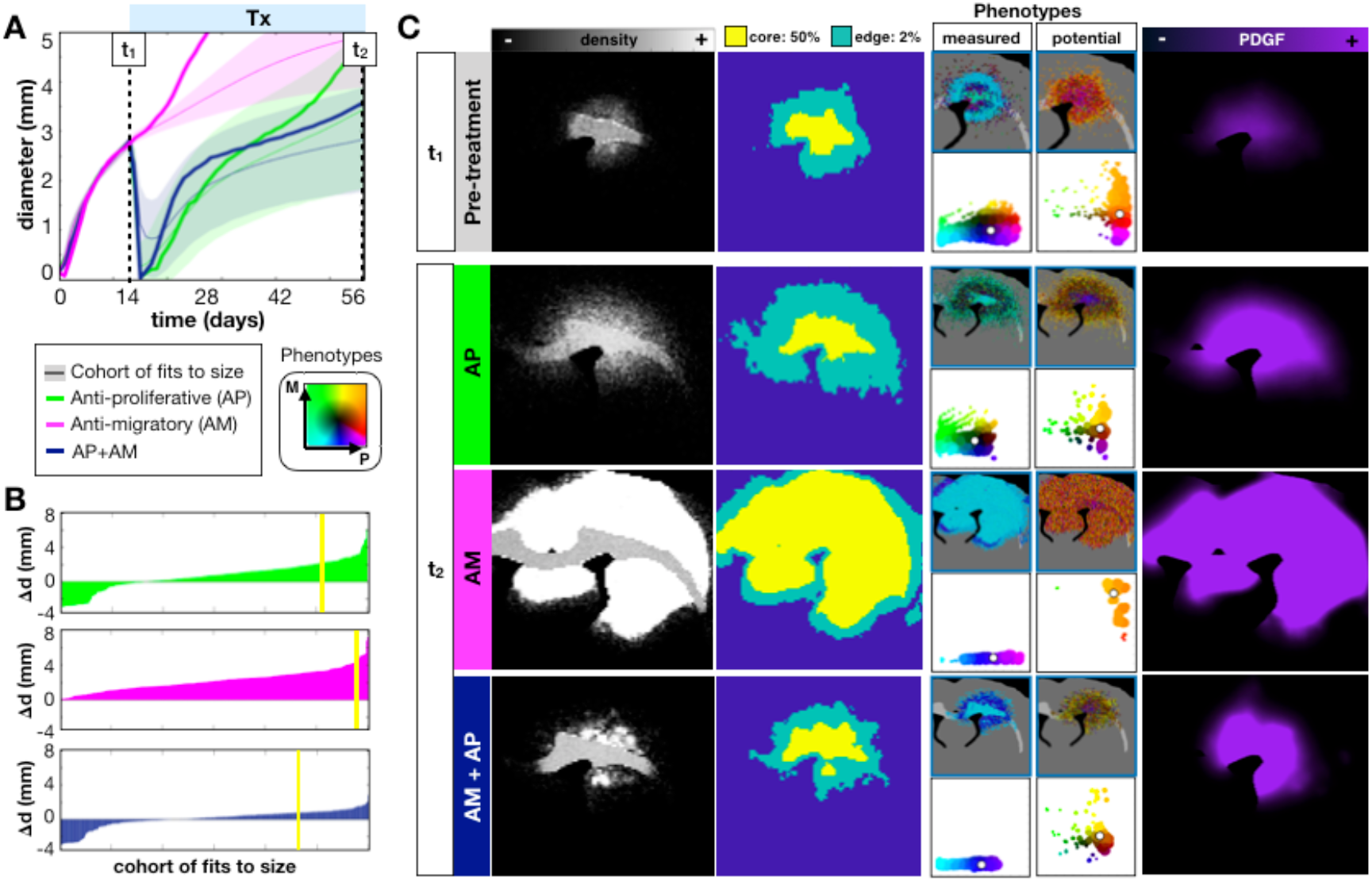
*In-silico* tumors do not respond well to an anti-migratory drug alone (AM), but may benefit from an anti-proliferative, anti-migratory (AP+AM) combination treatment. The drug is applied continuously at 14d until 28d. A) We show the growth dynamics for the AP, AM, and AP+AM treatments for the top 300 fits to the size dynamics. The average response (from 10 runs) to each treatment of the same diffuse tumor from the previous sections is also shown. B) Waterfall plot of the changes in tumor diameter from t_1_ to t_2_ for the cohort of top 300 fits to size when treated with AP (top) AM (middle), and AP+AM (bottom) treatments. The response of the diffuse tumor to these treatments is shown as a yellow line. C) Treating just the diffuse tumor example, we show the spatial density distributions, the core (>50% density) vs. the edge (>2% density), the measured and potential phenotype distributions (colored according to the key), and the PDGF distribution. The plots for the phenotype distributions are represented the same as described in Fig. 5.

The full cohort of *in silico* tumors fit to the size dynamics was examined for their response to the different treatments in Fig. 9B. We plot the change in tumor diameter before and after each treatment and see that a reduction in diameter in observed with 27% under AP, 0% with AM, and 36% with AP+AM. However, only 8% actually showed a complete response with either AP or AP+AM. Although in most cases, AP+AM resulted in better or similar outcomes than with AP alone (Fig. S11), in some cases, such as the representative diffuse tumor, a better response was seen with AP alone.

The response of diffuse tumor to each treatment is further examined in Fig. 9C. Prior to treatment, the tumor had a mean core diameter (d_c_) of 1.5mm with a mean rim size (d_r_) of 1.4 mm. With the AP treatment alone, the tumor appears to stay smaller for longer after treatment, but this measurement ignores many cells that invaded deep into the brain tissue under the imaging density threshold (at 41d, d_c_=2.6mm d_r_=2.8mm). With the AP treatment cells continue to migrate into the tissue, and slower proliferating cells are selected. With the AM treatment, the tumor grows very large since there is no killing taking place, but since the migration has essentially been turned off, growth is driven by proliferation alone rather than proliferation and dispersion (at 41d, d_c_=6.3mm d_r_=0.5mm). AM treatment selects for cells with high proliferative and migratory potential since they were previously selected for during growth and already populate the outer edges when migration is shut off. The PDGF concentration also becomes saturated in the tissue mediated by lack of cell dispersal, which further drives tumor growth. With the AP+AM treatment, the tumor is observed to be about the same size as the AP treatment (at 41d, d_c_=2.3mm d_r_=1.3mm), but the tumor was more cohesive and less diffuse. There was selection, again, for less proliferative cells after the AP+AM treatment, but the PDGF concentration was saturated within the tumor core. While the AP+AM treatment worked well over this time period for this tumor, a balance that needs to be made between preventing the widespread distribution of cells into the brain tissue and preventing the buildup of growth factor concentrations to such saturated levels that causes aggressive cell proliferation at the tumor core.

## Discussion

Tumor heterogeneity is fundamental to treatment success or failure. When predicting a tumor’s long-term response to treatment (observed on serial clinical imaging such as MRI), it is imperative to consider not just the change in the tumor size but also the variation in single cell phenotypes and heterogeneity in the environment. Our results suggest that growth rates alone are not enough to predict drug response; the tumor shape, density, and phenotypic and genotypic compositions can all signify characteristics of the underlying dynamics that affect longer term responses to therapy.

A tumor’s environmental context can play a huge role in malignant progression (5,38). We found through experiment and simulation that phenotypic heterogeneity is highly modulated by the environmental context. The local environment creates larger scale variations in the observed phenotypes that might be inhibiting, from factors such as lack of space or resources caused by a high cell density, or stimulatory, such as an overabundance of growth factors. These large-scale variations can give insight on environmental niches formed throughout the tumor. At the imaging scale, spatial variations can be quantified to reveal habitats and predict treatment response. Radiomic imaging does just that, because nuances in the shape, morphology, and texture of tumor density maps gives more information than size dynamics alone (3,6–8,18).

### Knowledge of intratumoral heterogeneity is required to predict patterns of treatment response and recurrence

Our results suggest that tumor heterogeneity is also not strictly a factor determined by the microenvironment, but a combination of cell autonomous drivers and the environmental context. *In silico* tumors that were fit to the same growth dynamics with similar density distributions displayed a huge variation in underlying phenotypes (Fig. 4). Furthermore, measurements at the single cell level do not necessarily match up with the potential behavior that cells could achieve given a different environmental context. It is often only after big changes in the tumor microenvironment, such as during therapy, that intrinsic variations at the single cell scale become apparent through natural selection (Fig. 5). Importantly, our data suggest that more information on single cell heterogeneity before treatment can lead to better treatment decisions. By fitting the *in silico* model to all of the experimental data, from bulk to single cell metrics, we found a best fit parameter set that resulted in a tumor with heterogeneity in the proliferative and migratory potential (Fig. 6). The best fit responded to an anti-proliferative drug but ultimately resulted in recurrence (Fig. 7). Eliminating the potential phenotypic heterogeneity in the best fit tumor did not drastically alter the resulting growth dynamics, yet upon exposure to the anti-proliferative treatment there was a complete response. Only at the single cell scale level (Fig. 6E) were we able to distinguish these two tumors that ultimately had divergent fates. From this result, it is clear that some degree of single cell observation could aid in the prediction of recurrence and a possible alteration of treatment strategy.

### Model prediction for response to anti-proliferative treatment is recapitulated in human patients

Based on our mathematical modeling results suggesting a diversity of phenotypes in response to treatment, we carefully investigated the role of anti-proliferative treatments since they form the basis of the vast majority of traditional anti-cancer treatments (e.g. radiation and chemotherapies). When fitting the mathematical model to the cell level and tissue level data, we found a consistent pattern of decreased proliferation in simulated recurrent tumors. This finding was recapitulated when we compared a histological marker for proliferation in human GBM patients at diagnosis and recurrence following chemoradiation (Fig. 8).

### Model predicts anti-migratory therapy may have limited impact as a monotherapy

Due to the invasiveness of GBM, the use of anti-migratory drugs is appealing (61,63–66). However, the *in silico* model suggests that anti-migratory drugs do not help when the tumor is largely driven by environmental factors (Fig. 9). Moreover, stopping migration also prevented the widespread dispersal of PDGF, leading to more proliferative tumors due to local accumulation of PDGF. This result indicates that, for this type of tumor, anti-migratory therapy alone is not significantly helpful. However, under the right conditions, it might be useful in combination with an anti-proliferative treatment or as a primer for an anti-proliferative drug. The anti-migratory drug was seen to select for more proliferative cells, so perhaps it could be used prior to an anti-proliferative treatment to select for more sensitive cells. Combining these treatments with an anti-PDGF drug could also help, to stop the response to environmental driving force in the first place (67).

### Model design limits interpretation of other biological mechanisms

In our model system we focused on phenotypic heterogeneity within a population of individual cells, which are modulated by the environment through cell density variation, the white/gray matter environment, and PDGF gradients. In order to simplify an already complex model that focuses on the relationship between cell autonomous heterogeneity and environmentally driven heterogeneity due to the growth factor, we excluded some significant drivers of environmental variation such as the angiogenic response, hypoxia, and necrosis (5,68,69). These are important components in the formation and progression of GBM in particular, however, in order to fit to the experimental data, we assumed that these factors played a backseat compared to the driving force of PDGF. While the *ex vivo* model allows for collection of data on multiple scales, it also represents an extreme case compared to human glioblastoma. The PDGF-driven rat model grows incredibly fast and recruits a large portion of resident progenitor cells by paracrine growth factor stimulation. The most sensitive parameter in the *in silico* model was the consumption rate of PDGF, which was quickly pushed to low values by the parameter estimation algorithm, a necessary component to promote a rapidly growing tumor. In the context of a slower growing tumor with less progenitor recruitment, which might be more accurate of human GBM, the growth, distribution, and evolution of cells could be quite different.

### A proliferation-migration dichotomy was not observed in the experimental data

We also made assumptions on the available phenotypes in this model, focusing on the most apparently important traits in GBM: proliferation rate and migration speed. A number of models and experiments find a limit to achieving both fast proliferating and fast migrating phenotypes, the idea of go-or-grow (70,71). However, even though it makes sense from a limited resource standpoint that cells have to divert energy from one task to another we found no dichotomy in the experimental data to warrant this assumption and perhaps an environment rich in growth factors caused no tradeoff. However, we found that *in silico* tumors with the same size dynamics tended to have measured proliferation and migration values that were not often both simultaneously high. It is possible that the proliferation-migration dichotomy is actually a consequence of environmental variation rather than a cell autonomous feature as seen in the model of Scribner *et al* (37). We also did not consider the impact of phenotypic evolution (13,41). The *ex vivo* data showed that the recruited cells, driven at least initially by the environment, proliferate and migrate faster than infected cells, which was found in the fully fit *in silico* model, but the rate of proliferation and migration of progenitor cells also increase over time. This could not be reiterated in the *in silico* model like the rest of the observations quantified here. If we were to consider phenotypic drift or transformation in the progenitor population, which has been reported to occur in other PDGF-driven glioma models (72), it is possible that the model would have fit the data better.

### Model suggests knowledge of intratumoral heterogeneity is required to effectively predict response to treatment

The *in silico* model allowed us to explore spatial dynamics of a tumor as a population and as individual cells to track heterogeneity over time and match to the experimental model. It showed that there likely needs to be both environmental and cell autonomous heterogeneity in order to fit to the smaller scale data, but these components are difficult if not impossible to separate by observation alone in a clinical setting. Specifically, there is no easy way to disentangle the drivers of observed phenotypic behaviors, since intrinsic cell autonomous drivers are modified by cell extrinsic environmental signals that themselves are modified by the cells. Here we have attempted to tackle this question through an integrated approach and hopefully shed light on this complex feedback. Using the hybrid agent-based model, we were able to combine data at different scales to study the environment and phenotypic heterogeneity separately and observe how single cell behavior influenced measurements at different scales. Although the anti-proliferative treatments showed variable responses in the *in silico* model, most were not sustaining and resulted in recurrence with slower proliferating, drug resistant phenotypes. Smarter strategies can be employed when more information is known about the tumor heterogeneity on all scales.

## Acknowledgments

We would like to thank the McDonnell Foundation for their support of this project (Collaborative Activity Award #220020264).

## Supplement

### S1. Single cell analysis

From the single cell tracks we quantified the migration behavior as depicted in Fig. S1. The speed for each cell over the time period was calculated as the total distance travelled over the total time spent moving. Since the cells frequently stopped for long periods of time, we excluded this from the calculation. Due to uncertainty in the cell’s center we defined a stopped cell as moving less than 5 *μ*m over the sampling time. The turning angles and persistence times were calculated by defining run times punctuated by frequent stops. We defined a single run as i) traveling a distance greater than 5 microns during the sampling time, and ii) continuing in the same direction to within 15 degrees of the original trajectory. A single stop time is just the amount of time spent before moving more than 5 *μ*m. The sampling frequency matters when capturing the observed speed and angle distributions (73,74). Due to the noisy data, which was recorded every 3 minutes, we sampled in 30-minute intervals, starting at different initial 3-minute time point within the 30-minute time interval, so no data was missed. Using these rules, we calculated turning angles, and persistence times.

**Figure S1.**
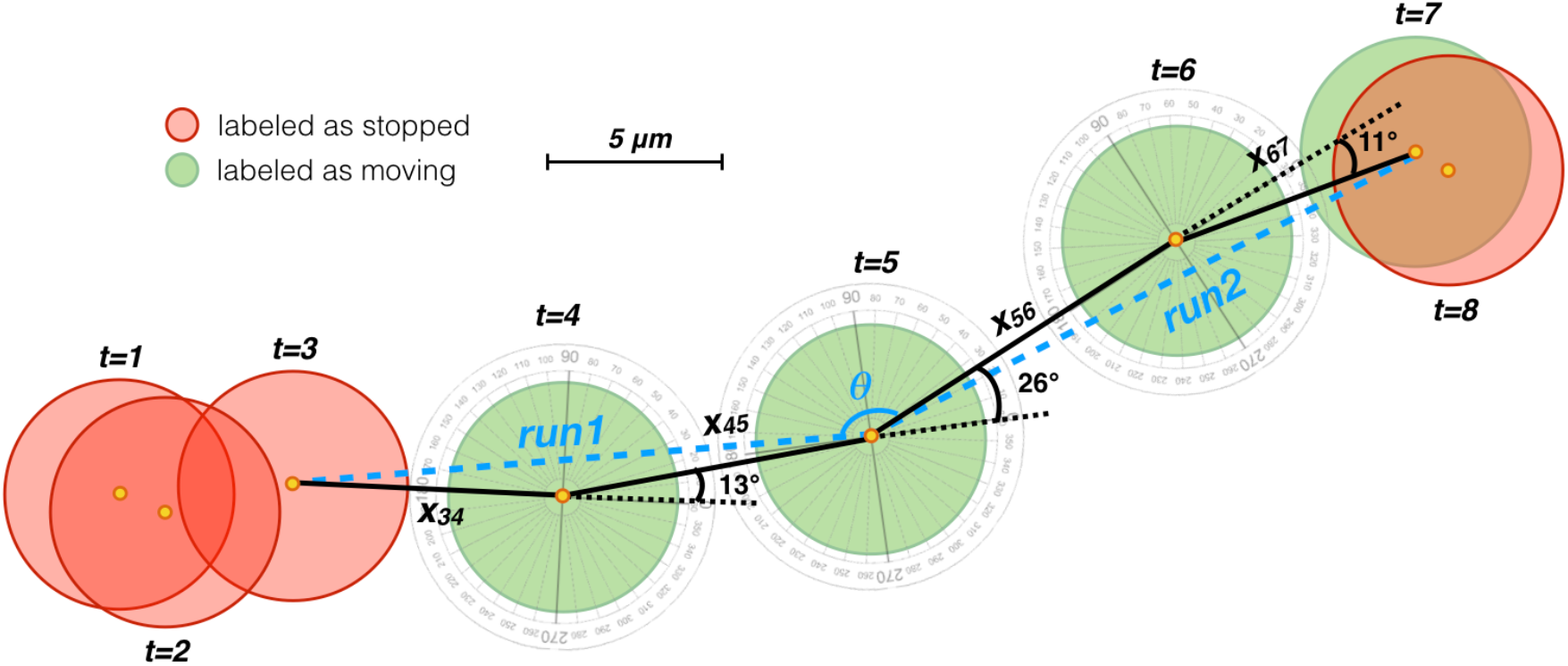
The cell track data analysis algorithm. Stops, runs, and turning angles are defined from each cell’s 2D track. The cell speed was calculated from the total distance travelled over the total time travelled (trajectory from solid black lines). If the cell moves greater than 5 *μ*m and does not turn more than 15 degrees during its trajectory, it is considered a single run (two runs and one turning angle labeled in blue).

### S2. Hexagonal lattice diffusion

We create a hexagonal lattice to store information on the type of tissue (gray or white matter) and the concentration of PDGF. Since the lattice has a staggered layout, it is indexed according to the scheme shown in Fig. S2. The concentration of PDGF at any time point is determined by adding to the concentration at the previous time point the sum of the differences between all the neighboring lattice sites times the diffusion coefficient and the time step over the distance travelled from the center of one lattice site to its neighbor. For each lattice point *i*, we write the concentration of PDGF as:

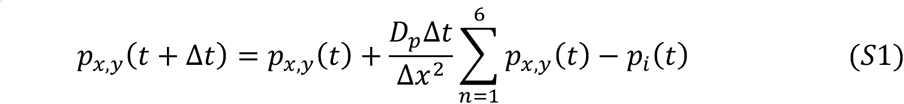

where is Δ*t* the time step, and Δ*x* is the distance from one lattice point to a neighboring one, which is just twice the apothem, so 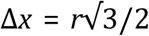 where *r* is the radius of the hexagon, which in this model is also the same size as the radius of a cell. There is no flux at the boundaries, so there is no contribution from off-grid neighbors.

**Figure S2.**
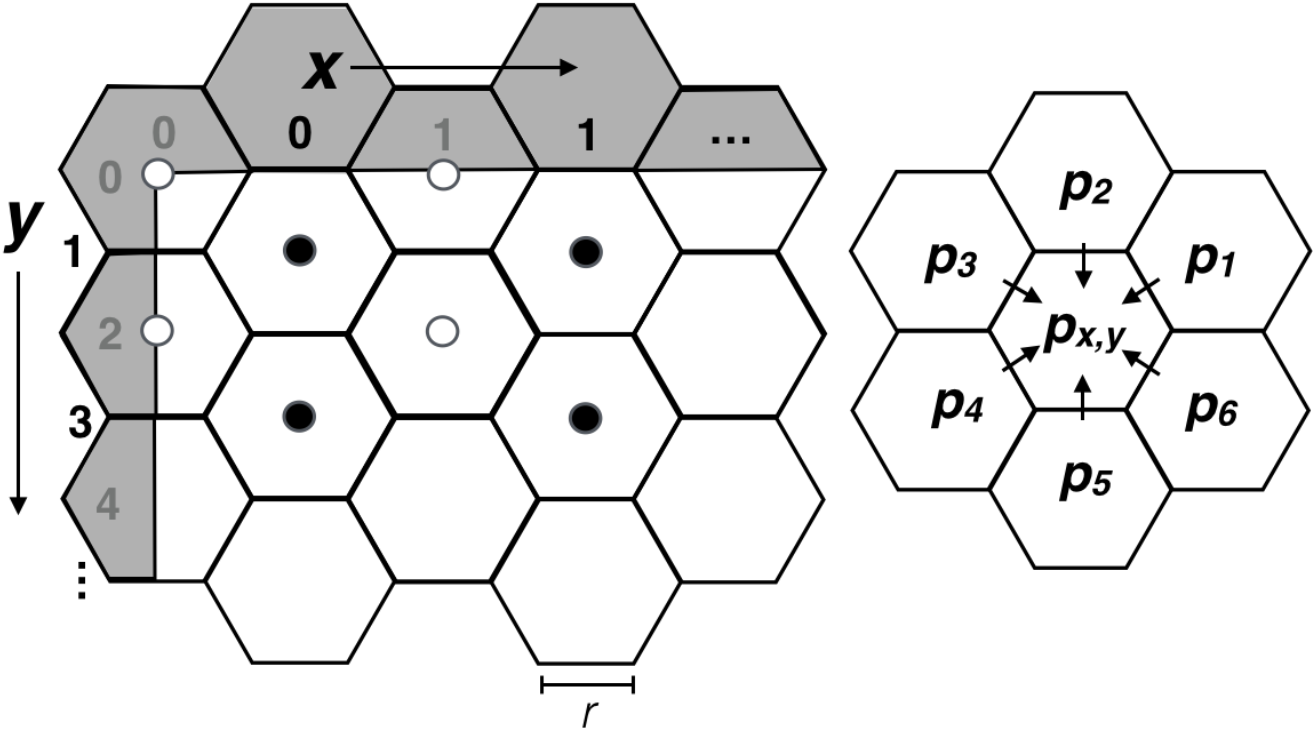
Hexagonal lattice diffusion. The lattice points are indexed as shown, with the even columns (black center dots) shifted down halfway in between the even rows (white center dots). Diffusion occurs at each lattice point p_x,y_ between the nearest neighbors only within the boundary of the domain. Off-grid lattice points (gray region) do not contribute to any flux.

### S3. Single cell data analysis

Using the detailed spatial movement behavior analysis explained in Section 1, we analyzed the data to learn more about migration patterns. The results are shown in Fig. S3 and described in detail here.

**Figure S3.**
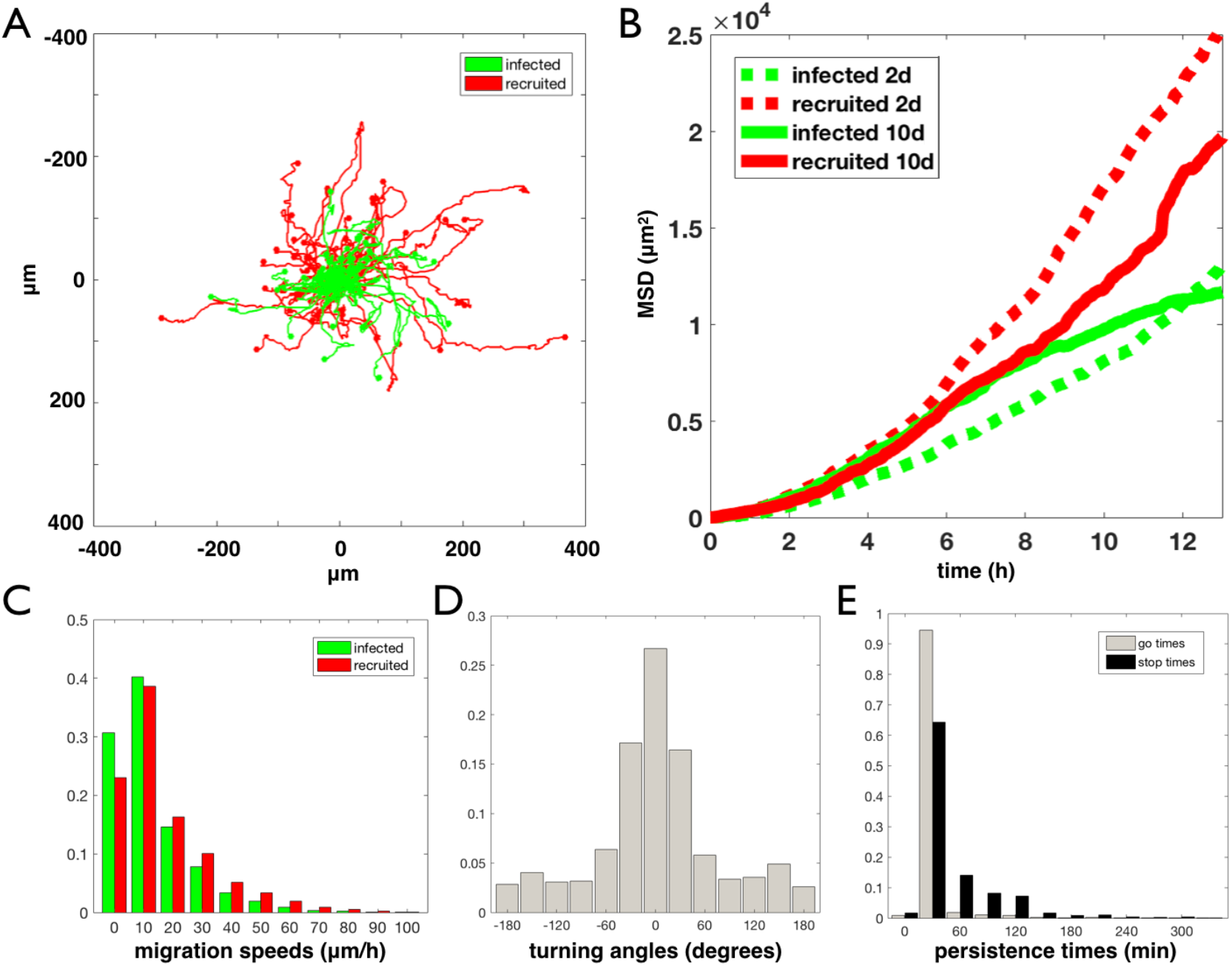
Behavior of single cells from rat data. A) Wind-Rose plot for infected and progenitor cells at 10d, B) Mean squared distance (MSD) for infected and recruited cells at both 2d and 10d, C) distribution of migration speeds at 10d, D) distribution of turning angles averaged over infected and recruited cells at 10d, and E) distribution of moving and stopping persistence times at 10d averaged over all cells.

#### Cell trajectories and distribution

To visualize the general distribution of cells over the time period of the movie, we show a Wind Rose plot of the trajectories (Fig. S3A). This plot shows the trajectories of all cells in one slice of data at 10d with all of the starting points for each cell placed at the same origin. It is observed that the cells have a variety of paths with a variety of distances travelled. The red progenitor cells appear to move farther over the same time period.

#### Mean squared distances

To quantify how far the population of cells moved over the length of time of the movie, we calculated the mean squared distances (MSDs) for each population at each time point (Fig. S3-B). At both time points, the red progenitor cells are seen to have a greater MSD slope. However, while the green infected cells have similar slopes at both time points, the red progenitor cells appear to slow down at the later time point.

#### Migration speeds

We calculated the migration speeds for each cell using the method in Section S1 and found that there is a large variation in the cell speeds, going up to 100 μm/h but peaking round 10 μm/h (Fig. S3-C). Because the both time points had similar distributions, we grouped the two time points into one plot, but the distribution of progenitor cells speeds is slightly shifted toward higher values.

#### Turning angles

The turning angles are calculated to have a narrow peak at 0 degrees, indicating that there is a lot of persistent movement, but there is also a broad uniform coverage of larger angles (see Fig. S3-D).

#### Persistence times

The cells move and stop often over the 25-hour time period of the movie. We found that the overall time spent moving and not moving is not significantly different between infected and progenitor cells. However, on average, cells are more often stopped than moving (Fig. S4-E).

#### Values for model fitting

The values gathered from the data from which we matched to the model are summarized in Table S1. This contains tumor scale data from imaging, and single cell scale data from the tissue slice data.

**Table S1.** Data measured from the rat experiment that was used to fit the model. The larger scale data was taken from Assanah et al (12) and Massey et al (44), whilst the single cell data was calculated from the cell tracks as stated in the methods.

| PARAMETER | SCALE | TIME POINT (d) | VALUE | SOURCES |
| --- | --- | --- | --- | --- |
| Diameter (mm) | tumor | 5 | 1.7 | (14,33) |
|  | tumor | 10 | 2.4 | (14,33) |
|  | tumor | 17 | 3.2 | (14,33) |
| Ratio I/R | tumor | 17 | 0.2 | (14) |
| mean proliferation rate<br>(% cells/h) | infected cells | 2 | 0.33 | calculated |
|  | recruited cells | 2 | 0.85 | calculated |
|  | infected cells | 10 | 0.83 | calculated |
|  | recruited cells | 10 | 1.89 | calculated |
| mean migration rate<br>($\mu\text{m}/\text{h}$) | infected cells | 2 | 21.3 | calculated |
|  | recruited cells | 2 | 24.9 | calculated |
|  | infected cells | 10 | 20.6 | calculated |
|  | recruited cells | 10 | 25.2 | calculated |
| standard deviation<br>migration ($\mu\text{m}/\text{h}$) | infected cells | 2 | 5.7 | calculated |
|  | recruited cells | 2 | 7.6 | calculated |
|  | infected cells | 10 | 5.7 | calculated |
|  | recruited cells | 10 | 8.8 | calculated |

### S4. Parameter estimation

#### Matching model to data metrics

##### Tumor size

Since the tumor consists of cells often spread out with no clear boundary, the tumor size was measured by finding the maximum distance from the center of mass where the cell density is >= 10% of the carrying capacity (averaged over 12 angles).

##### I/R

The ratio of infected to recruited cells is calculated, as with the data, by observing the number of each cell type within the tumor core.

##### Single cell metrics

For the single cell metrics, a randomly chosen subset of up to 200 cells were tracked of the initially injected and labeled subset (as opposed to native inactive progenitors) outside of the densest regions (less than 25% of the carrying capacity). These criteria bias the tracks to the tumor edge where it is not too dense to agree with the experimental tracking limitations. Proliferation events and positions of the cells were tracked every 3 min. The measured proliferation rate and migration speeds were calculated by recording the percent of divisions over the observation period and the total distance traveled over the time spent moving, respectively. These values were averaged for infected and recruited cells.

### Convergence scheme

There are 16 free parameters that are either a) not measured, b) not well determined by experimental estimates, or c) variables in the simulation. These are listed in Table 1 of the main text.

To converge on reasonable parameter estimates that result in good fits to the data in Table S1, we start by drawing 5000 sets of random values for each parameter within the determined ranges. From each parameter set we run the simulation, calculate the output values for each metric, and compare to the data output to get a total error. We then sort these from least to most error and take the top 10%, excluding any set of parameters that have more than 50% of the total error in one metric, and transfer these directly to the next iteration. For the next 40% of the next iteration, we tweak all of the parameters in the top 10% by a random value sampled from a normal distribution with a standard deviation of 10% of the parameter range. For the final 50% of the next parameter set, we draw randomly from the parameter distribution of the top 10% to introduce new combinations of values. We iterate this procedure until the output error is within 10% for fitting the size dynamics, which turns out to be after 5 iterations, and within 15% for fitting all metrics, which is after 13 iterations. The distributions at each iteration are shown in Fig. S4 and the final parameter values used for each tumor type are given in Table S2.

**Figure S4.**
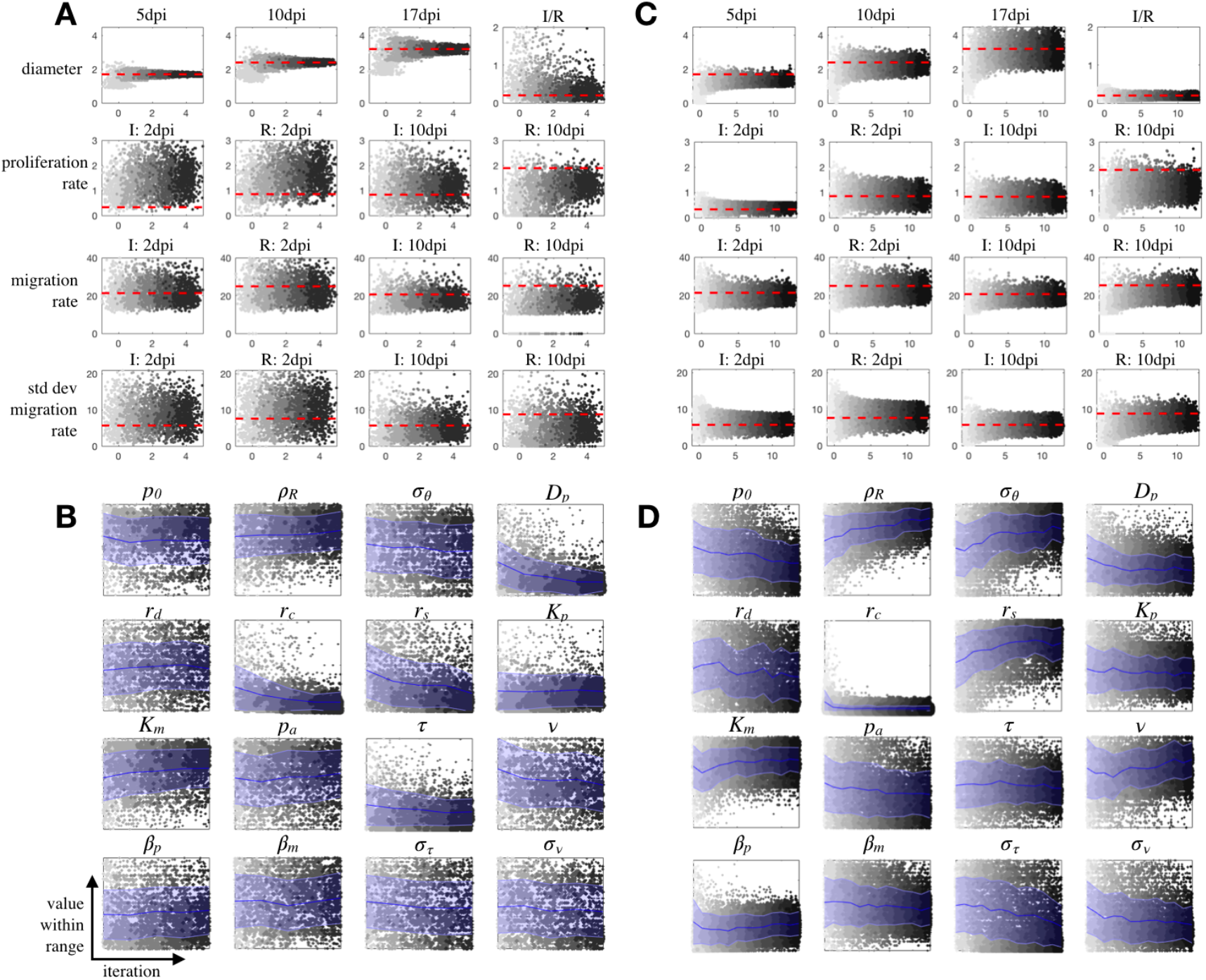
Parameter estimation by matching to data. Values over iterations of the convergence are shown for A) metrics of top 300 fits fit to size dynamics only, B) parameters from the top 300 fits to size dynamics only, C) metrics of top 300 fits using all data, and D) parameters from the top 300 fits using all data. Each iteration is shown starting at light gray and going to black for the final fit. The red dashed line for the metrics indicates the measured data values, while the blue lines and error bars show the mean and standard deviation over iterations for each parameter.

**Table S2.**
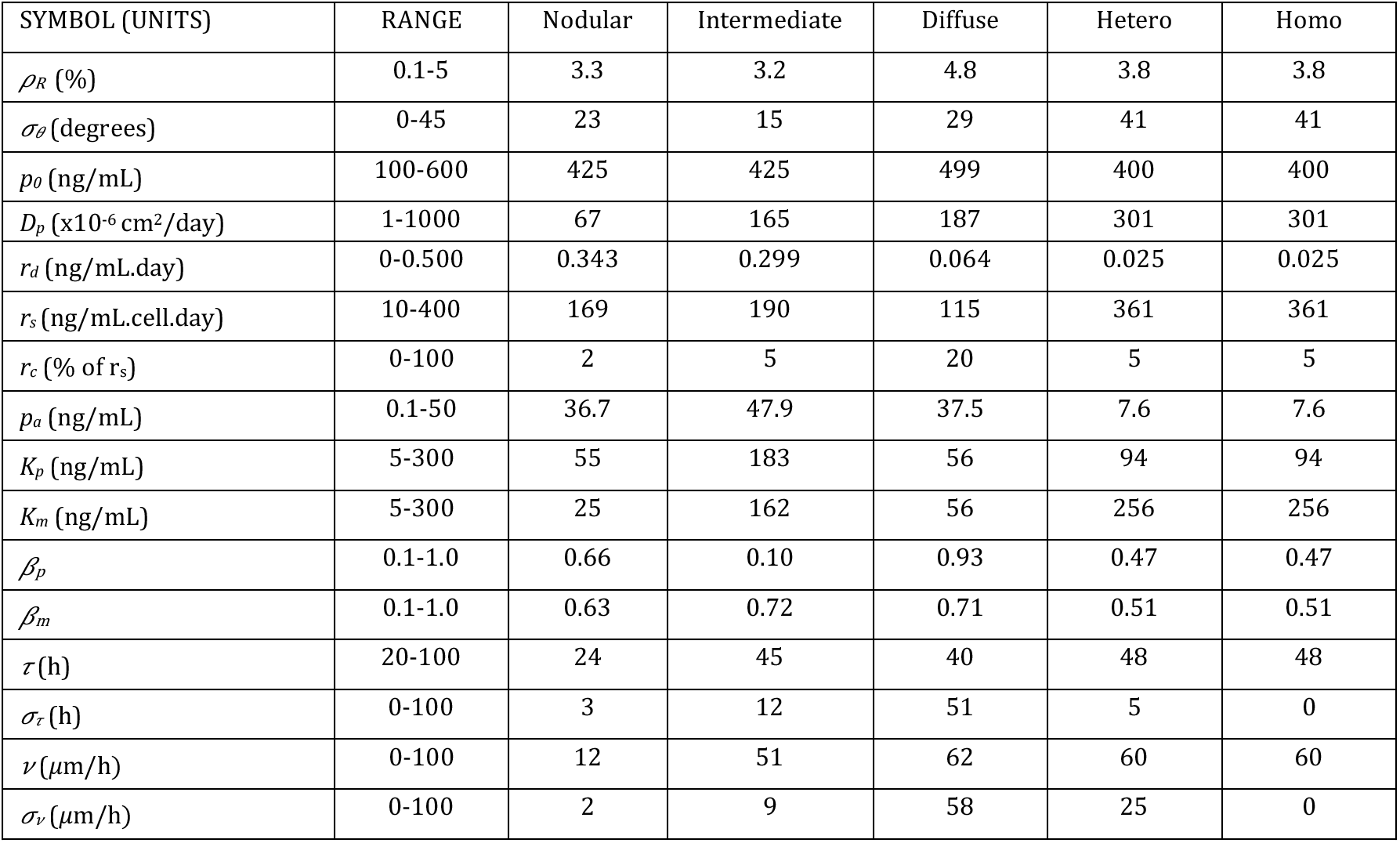
Parameter sets used for the example tumors in main text. The parameter ranges are used to search for fits to the data. The nodular, intermediate, and diffuse tumors are found by fitting only to the tumor size data, and the heterogeneous tumor is found by fitting to all of the data. The homogeneous tumor is just the heterogeneous tumor with the variation in proliferation and migration set to zero.

### S5. Supplemental Results

**Figure S5.**
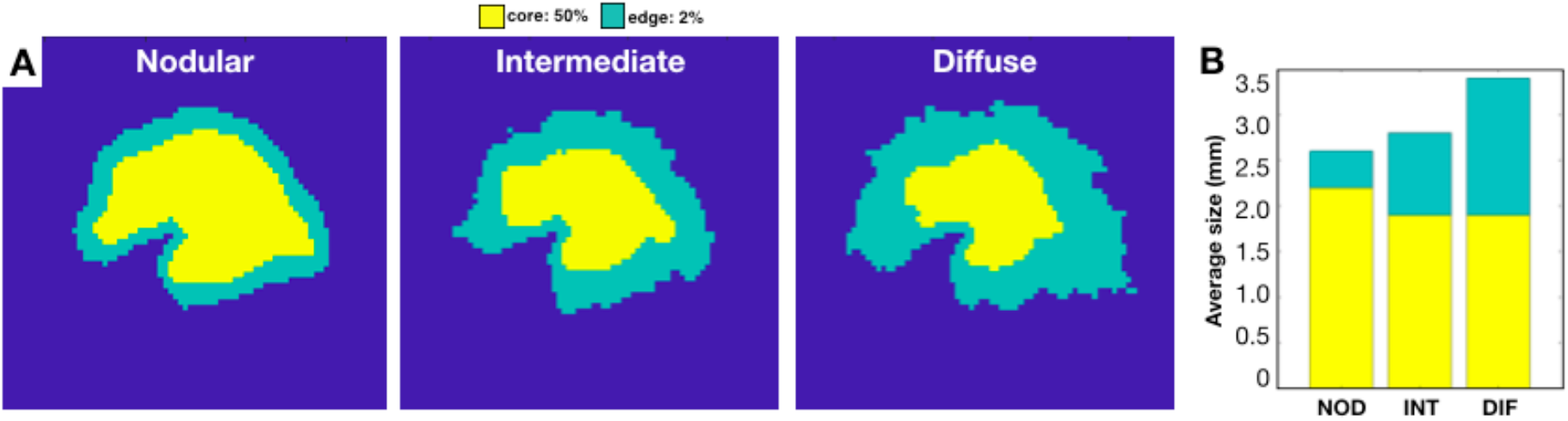
Tumor density profiles. A) For the nodular, intermediate, and diffuse tumors, the core (yellow) is defined as having a cell density of at least 50% of the carrying capacity, while the rim (green) is defined as having a cell density of at least 2% of the carrying capacity. B) Stacked bar plot of average core diameter and average rim diameter over 10 runs. We define the average rim size as the difference between the average rim diameter and the average core diameter. The average core diameters were 2.2mm, 1.9mm and 1.9mm for the nodular, intermediate, and diffuse tumors, and the average rim sizes were 0.4mm, 0.9mm, and 1.5mm, respectively.

**Figure S6.**
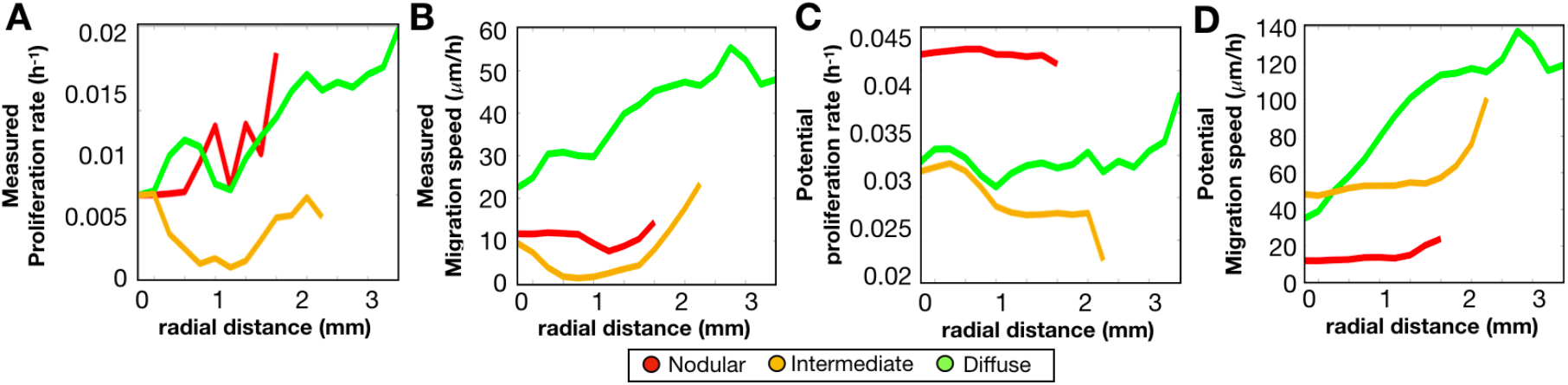
Spatial phenotype distributions along the radius of the tumor are shown at 17d. The average values over 10 runs are plotted: A) measured proliferation rate, B) measured migration speed, C) potential proliferation rate, and D) potential migration speed.

**Figure S7.**
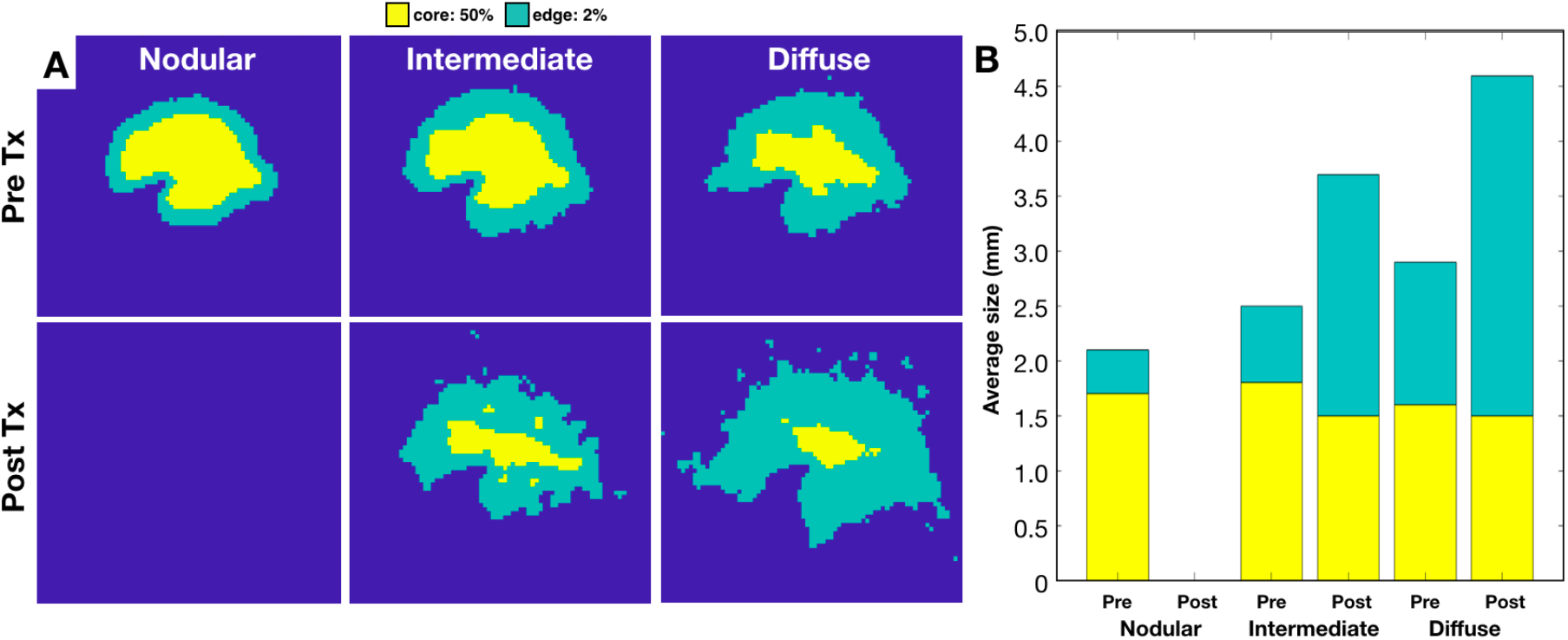
Changes in density profiles of nodular, intermediate, and diffuse *in-silico* tumors before and after an anti-proliferative treatment application. A) For the nodular, intermediate, and diffuse tumors, the core and rim as defined in Fig. S5 are shown. B) Stacked bar plot of average core diameter and average rim diameter over 10 runs. The average core diameter pre-treatment was 1.7mm, 1.8mm and 1.6mm for the nodular, intermediate, and diffuse tumors, and post-treatment were 0mm, 1.5mm, and 1.5mm, respectively. The average rim size pre-treatment was 0.4mm, 0.7mm, and 1.3mm for the nodular, intermediate, and diffuse tumors, and post-treatment were 0mm, 2.2mm, and 3.1mm, respectively.

**Figure S8.**
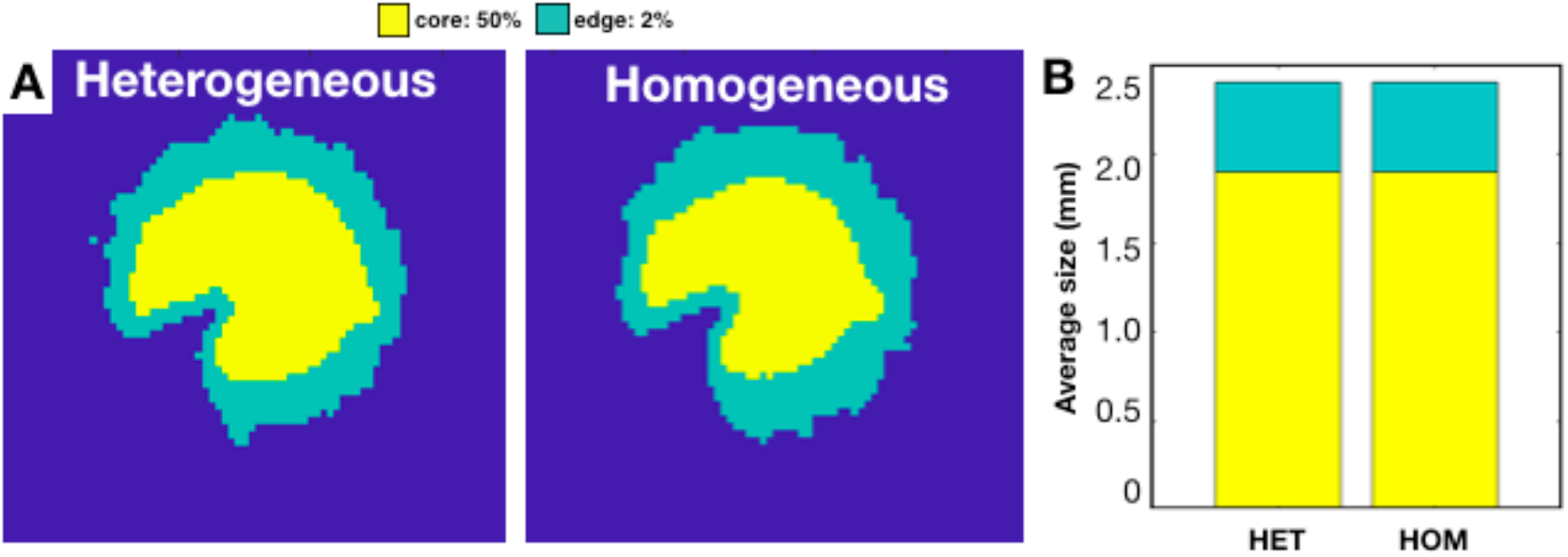
Density profiles of the heterogeneous and homogeneous *in-silico* tumors. A) The core and rim as defined in Fig. S5 are shown. B) Stacked bar plot of average core diameter and average rim diameter over 10 runs. The average core diameters were both 1.9mm, and the average rim sizes were both 0.5mm.

**Figure S9.**
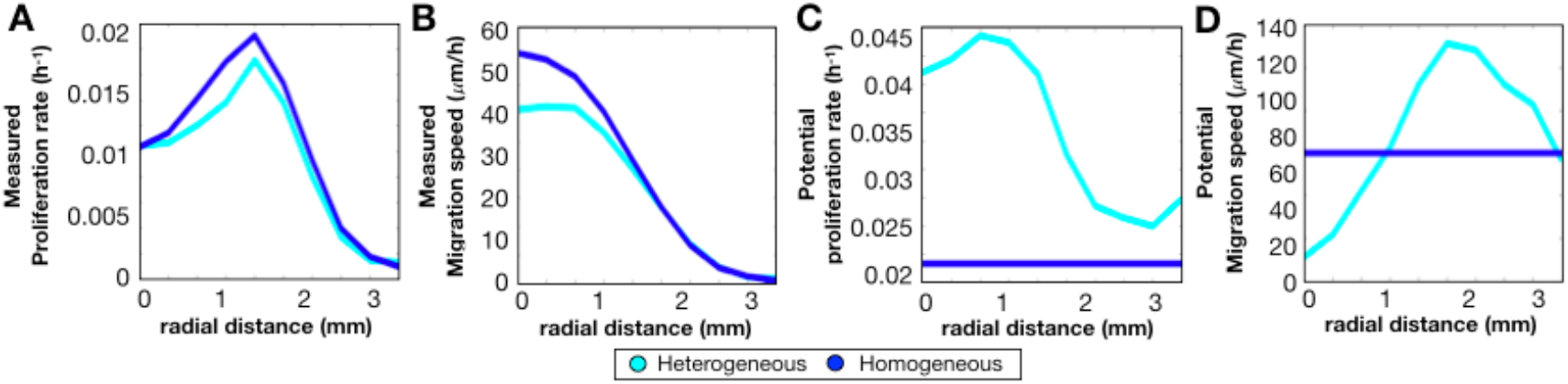
Spatial phenotype distributions along the radius of the tumor are shown at 17d. The average values over 10 runs are plotted: A) measured proliferation rate, B) measured migration speed, C) potential proliferation rate, and D) potential migration speed.

**Figure S10.**
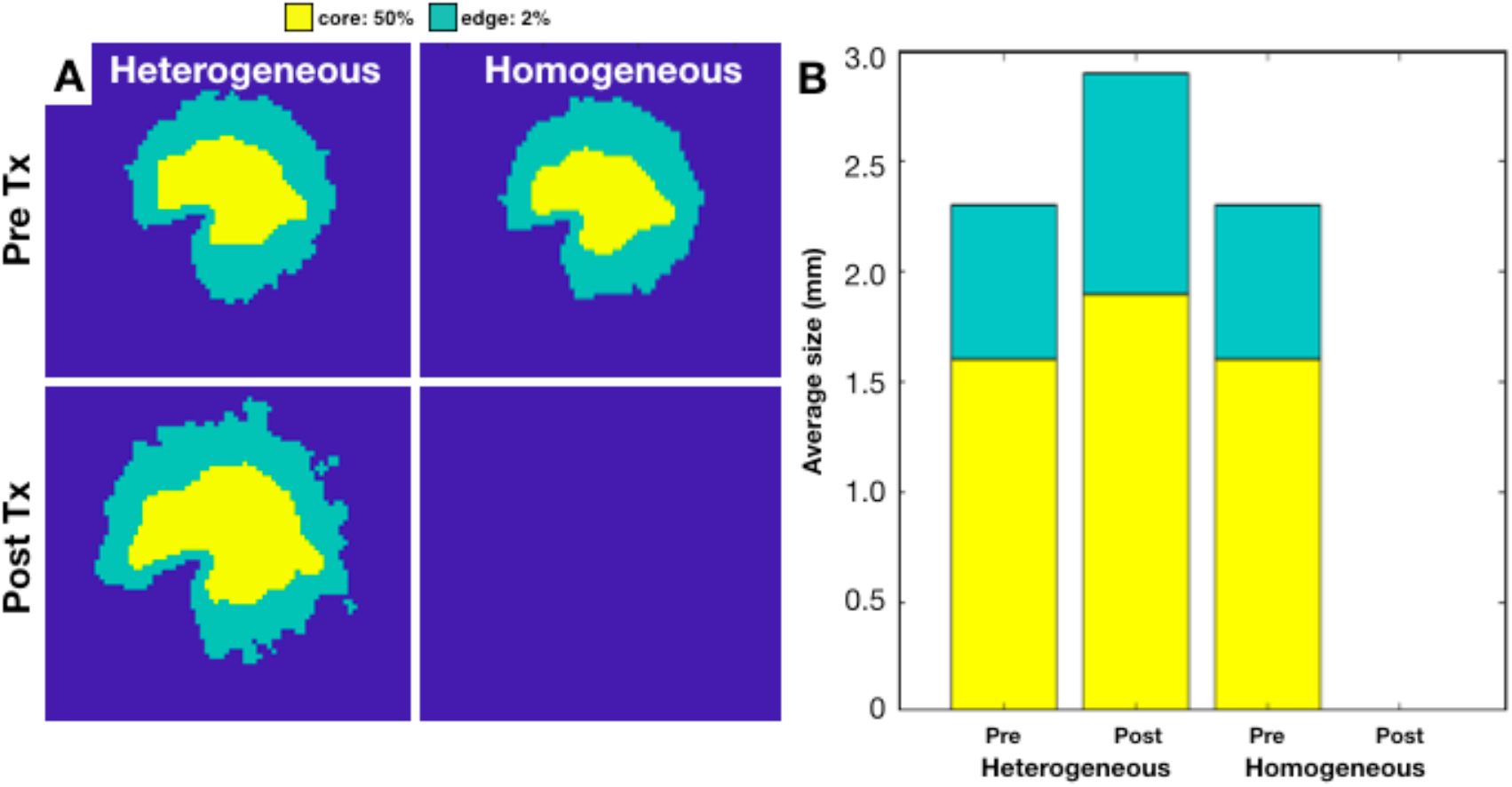
Changes in density profiles of the heterogeneous and homogeneous *in-silico* tumors before and after an anti-proliferative treatment application. A) The core and rim as defined in Fig. S5 are shown. B) Stacked bar plot of average core diameter and average rim diameter over 10 runs. The average core diameter pre-treatment was 1.6mm for both, and the post-treatment heterogeneous tumor was 1.9mm. The average rim size pre-treatment was 0.7mm for both, and the post-treatment heterogeneous tumor was 1.0mm.

**Figure S11.**
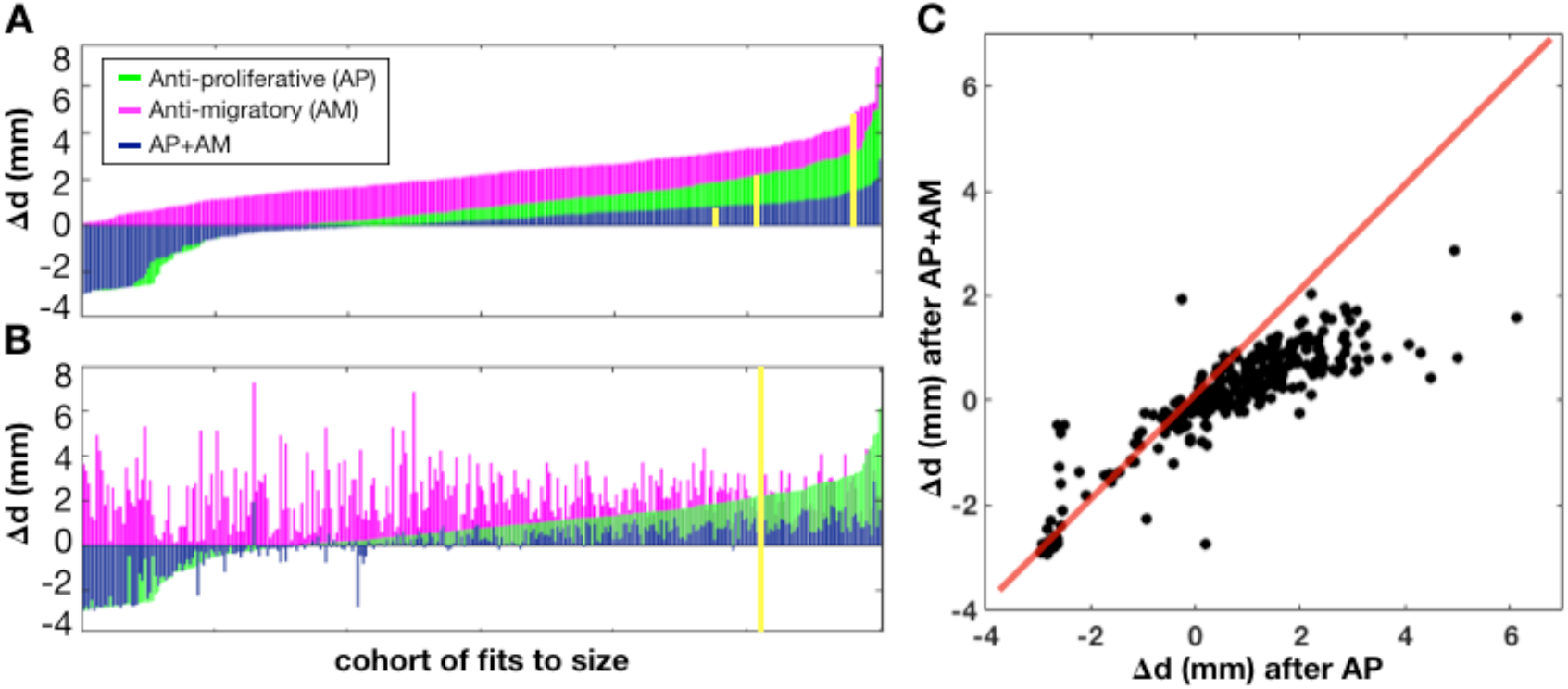
Correlation between treatment outcomes over cohort of simulated tumors. We show the distribution of response as A) a waterfall plot with each treatment sorted ranked from best to worst response and B) a waterfall plot for AP treatment sorted ranked from best to worst response but preserving the correlation of how each tumor responds to the other treatments. The yellow line shows the responses for the diffuse tumor from Fig. 9. C) Comparison of the responses for AP treatment alone to AP+AM combination treatment. The red line shows where the response is the same for both treatments.

## Notes

#### Summary of Updates

Fixed a typo and author affiliation.

